# Socio-behavioural characteristics and HIV: findings from a graphical modelling analysis of 29 sub-Saharan African countries

**DOI:** 10.1101/600510

**Authors:** Zofia Baranczuk, Janne Estill, Sara Blough, Sonja Meier, Aziza Merzouki, Marloes H. Maathuis, Olivia Keiser

## Abstract

**Introduction:** Socio-behavioural factors may contribute to the wide variance in HIV prevalence between and within sub-Saharan African (SSA) countries. We studied the associations between socio-behavioural variables potentially related to the risk of acquiring HIV.

**Methods:** We used Bayesian network models to study associations between socio-behavioural variables that may be related to HIV. A Bayesian network consists of nodes representing variables, and edges representing the conditional dependencies between variables. We analysed data from Demographic and Health Surveys conducted in 29 SSA countries between 2010 and 2016. We predefined and dichotomized twelve variables, including factors related to age, literacy, HIV knowledge, HIV testing, domestic violence, sexual activity, and women’s empowerment. We analysed data on men and women for each country separately and then summarised the results across the countries. We conducted a second analysis including also the individual HIV status in a subset of 23 countries where this information was available. We presented summary graphs showing associations that were present in at least six countries (five in the analysis with HIV status).

**Results:** We analysed data from 190,273 men (range across countries 2,295–17,359) and 420,198 women (6,621–38,948). The two variables with the highest total number of edges in the summary graphs were literacy and rural/urban location. Literacy was negatively associated with false beliefs about AIDS and, for women, early sexual initiation, in most countries. Literacy was also positively associated with ever being tested for HIV and the belief that women have the right to ask their husband to use condoms if he has a sexually transmitted infection. Rural location was positively associated with false beliefs about HIV and the belief that beating one’s wife is justified, and negatively associated with having been tested for HIV. In the analysis including HIV status, being HIV positive was associated with female-headed household, older age and rural location among women, and with no variables among men.

**Conclusions:** Literacy and urbanity were strongly associated with several factors that are important for HIV acquisition. Since literacy is one of the few variables that can be improved by interventions, this makes it a promising intervention target.

## Introduction

In sub-Saharan Africa, about 26 million people were living with HIV in 2018 [1]. HIV prevalence is high overall, but very heterogeneous between countries, ranging from under 1% in Senegal and Niger to up to 25% in Lesotho and eSwatini (previously Swaziland) [2–6]. The HIV prevalence also varies substantially within countries, the overall epidemic being concentrated in clustered micro-epidemics of different geographical scales [7]. The Joint United Nations Programme on HIV/AIDS (UNAIDS) has urged researchers to pay attention to these variations, and to identify geographical areas where the risk of acquiring HIV is higher: “know your epidemic, know your response” [7]. Variation has been attributed to many factors, including socio-demographic, behavioural, and biological [3,8]. Among the factors associated with HIV burden are age, religion, marital status, high levels of HIV-related stigma, sexual coercion, education, occupation, and gender [4,8].

Education can affect HIV prevalence. Even in countries where everyone knows HIV risk is high, the public may not know much about how the disease spreads. For example, those who have not been taught why and how to use condoms are less likely to protect themselves [8]. Attitudes and behaviour can also contribute to HIV risk. If false beliefs about HIV and AIDS are not corrected and replaced with specific, accurate knowledge, this can increase risky sexual behaviour and infection risk [9]. Women with little education are likely to be less informed about interventions that mitigate HIV risk [9], but higher HIV rates among women cannot be fully explained by either lower educational levels or higher biological susceptibility. In countries with high HIV prevalence, gender inequality has been associated with HIV spread along with low levels of general education [10] and sexual education [8]. Women with limited or no income may engage in transactional sex to survive [11], so socioeconomic and living conditions can also be factors in HIV transmission. Urban residence may increase the risk of infection by increasing the probability of high risk sexual contacts [12], but it may also decrease HIV risk because urban dwellers are more likely to be employed and have better access to care and treatment [13].

Most studies about risk factors for HIV or for the uptake of interventions against HIV have used logistic regression models to simultaneously adjust for a range of factors [14,15]. Such an approach, however, does typically not account for the complex network of associations and causal pathways between various risk factors and HIV. Understanding this network could help us identify drivers of the epidemic to plan effective interventions and ultimately reach the global treatment targets set by United Nations Member States [2]. We thus explored the associations between socio-behavioural variables that could contribute to the HIV epidemic by analysing data from Demographic and Health surveys from 29 countries in Sub-Saharan Africa using Bayesian network analysis [16].

## Methods

### Data description

The Demographic and Health Surveys (DHS) Program collects and disseminates high-quality, nationally representative data on fertility, family planning, maternal and child health, gender, HIV/AIDS, malaria, and nutrition in low- and middle-income countries around the world. Typically, it collects data on between 3,300 and 5,300 variables per country every 5 years. The DHS surveys include women of reproductive age (15-49 years) and (depending on the country) men aged 15-49, 15-54 or 15-59 years.

We used data from the DHS and included all countries in sub-Saharan Africa where a DHS survey was administered during or after 2010 [6]. We included the last available survey for each country (as of August 2018): Angola 2015-16; Benin 2011-12; Burkina Faso 2010; Burundi 2010; Cameroon 2011; Chad 2014-15; Congo 2011-12; Democratic Republic of the Congo 2013-14; Côte d’Ivoire 2011-12; Ethiopia 2016; Gabon 2012; The Gambia 2013; Ghana 2014; Kenya 2014; Lesotho 2014; Liberia 2013; Malawi 2015-16; Mali 2012-13; Mozambique 2011; Namibia 2013; Niger 2012; Nigeria 2013; Rwanda 2014-15; Senegal 2010-11; Sierra Leone 2013; Togo 2013-14; Uganda 2011; Zambia 2013-14; and, Zimbabwe 2015. We excluded Tanzania from the analysis because some essential variables were missing from their DHS data.

Two researchers (ZB and OK) preselected variables and dichotomized them before analysis. We included the following variables because they covered topics that could relate to HIV and were available for all selected countries: age 24 or younger (yes, no); rural location (yes, no); female head of household (yes, no); literacy (able to read whole sentence: yes, no/missing); access to media at least once a week (yes, no/missing); sexual initiation before 16 years (yes, no); currently working (yes, no/missing); judging wife-beating justified (yes/does not know/missing, no); married or living together either now or previously (yes, no); false beliefs about AIDS (yes/does not know/missing, no); judging it justified for a wife to ask her husband to use a condom if he has a sexually transmitted infection (STI; yes, no/does no know/missing); ever tested for HIV (yes, no/missing); and diagnosed HIV positive at the time of the survey (yes, no/no result) [17].

### Data analysis

We conducted exploratory data analyses and Bayesian network analyses separately for men and women in each country and included all variables except HIV status in our main analysis. The information on HIV status was available only for 23 countries, so we performed separate analyses that included HIV status for these countries.

First, we calculated and presented crude odds ratios for each pair of variables by country and sex. Then we conducted a Bayesian network analysis to determine the possible causal relationship between variables for each country, separately for men and women. Bayesian networks are graphical models in which nodes represent random variables and edges represent certain probabilistic dependencies between them. In particular, the presence of an edge means that the corresponding variables are dependent given any subset of the remaining variables. Our analysis assumes that there are no unmeasured confounders. While the presence or absence of an edge is always identifiable under our assumptions, the directions of edges may be unknown, so we can only learn a so-called completed partially directed acyclic graph (CPDAG) [18,19]. The theoretical framework for CPDAG learning is well-developed [20-23]. We used a fully automatic approach with the Incremental Association Markov Blanket (IAMB) algorithm [24] within the bnlearn package in R [25]. To improve the stability of the estimated graphs, we used bootstrap sampling. For each country and sex, we took 10,000 bootstrap samples, using weighted resampling to account for the sampling weights in the DHS data. For each bootstrapped data set, we obtained the corresponding estimated CPDAG. We then constructed a graph for each country as follows: two nodes were connected if the edge was present in at least 90% of the bootstrapped CPDAGs, and edges were oriented if they were oriented in that direction in at least 70% of the bootstrapped CPDAGs.

Finally, we determined an overall summary graph by sex that combined the information for all countries. In this summary graph we only considered edges present in at least six countries for the analysis without HIV status, or five countries for the analysis with HIV status. The orientation of edges in the summary graph was determined as follows: For each edge, the number of countries with an orientation in one direction, and the number of countries with an orientation in the other direction was determined. If the difference between these numbers of countries was at least six, the edge in the summary graph was oriented according to the majority orientation. We estimated the sign of an association conditionally on all other variables using the Cochran-Mantel-Haenszel odds ratio [26] and marginally by calculating the crude odds ratio. We considered associations negative if both the crude and the conditional odds ratio were less than one, and positive if both the crude and the conditional odds ratios were above one.

We used the open source R language, version 3.5.0 for our analysis [25]. The code is available on github (https://github.com/ZofiaBTZ/DAGS_DHS).

### Ethical approval

The study was conducted using data from the DHS program, which are freely available on request, and thus no ethical approval was needed. Details of the ethical review process of DHS are available on the programme’s website https://www.dhsprogram.com/What-We-Do/Protecting-the-Privacy-of-DHS-Survey-Respondents.cfm.

## Results

We included 190,273 men (range across countries 2,295–17,359) and 420,198 women (6,621– 38,948) in the analysis. The age distribution and percentage of people living in urban areas were similar for males and females (Table 1). Women were less likely to be able to read (median 46% vs. 68%), to have media access (61% vs. 74%), to currently work (54% vs. 79%), and more likely to have been tested for HIV (49% vs. 31%).

**Table 1.** Characteristics of persons included in the analysis.

| DHS, 2010 - 2016, countries of sub-Saharan Africa |  | Female<br>median (min, max) | Male<br>median (min, max) |
| --- | --- | --- | --- |
| Younger than 25<br>years | No | 60.0% (54.7%-65.5%) | 62.9% (56.1%-72.1%) |
|  | Yes | 40.0% (34.5%-45.3%) | 37.1% (27.9%-43.9%) |
| Rural setting | No | 38.7% (10.4%-88.5%) | 40.2% (14.9%-87.1%) |
|  | Yes | 61.3% (11.5%-89.6%) | 59.8% (12.9%-85.1%) |
| Household head<br>female | No | 72.7% (43.7%-90.8%) | 86.8% (71.8%-97.1%) |
|  | Yes | 27.3% (9.2%-56.3%) | 13.2% (2.9%-28.2%) |
| Literacy | No (Not able to read whole<br>sentence/missing) | 56.6% (9.6%-89.7%) | 32.0% (14.0%-74.1%) |
|  | Yes (Able to read whole sentence) | 43.4% (10.3%-90.4%) | 68.0% (25.9%-86.0%) |
| Media access | Less than once a week | 39.5% (8.0%-81.7%) | 25.7% (4.2%-64.8%) |
|  | At least once a week | 60.5% (18.3%-92.0%) | 74.3% (35.2%-95.8%) |
| First sex before the<br>age of 16 | No | 67.4% (50.5%-93.6%) | 79.9% (54.6%-97.7%) |
|  | Yes | 32.6% (6.4%-49.5%) | 20.1% (2.3%-45.4%) |
| Currently working | No | 46.5% (24.8%-79.1%) | 21.4% (6.9%-44.0%) |
|  | Currently working / have a job, but on<br>leave during the last 7 days | 53.5% (20.9%-75.2%) | 78.6% (56%-93.1%) |
| Married | Never in partnership | 28.3% (8.6%-53.5%) | 40.8% (28.7%-62.9%) |
|  | Currently or formerly married or living<br>with a partner | 71.7% (46.5%-91.4%) | 59.2% (37.1%-71.3%) |
| False beliefs about<br>AIDS | No | 55.5% (35.8%-88.4%) | 62.3% (44.5%-85.7%) |
|  | Yes / Does not know/ Missing | 44.5% (11.6%-64.2%) | 37.7% (14.3%-55.5%) |
| Wife beating<br>justified | No | 50.6% (20.6%-83.3%) | 88.5% (66.4%-95.5%) |
|  | Yes / Does not know/ Missing | 49.4% (16.7%-79.4%) | 11.5% (4.5%-33.6%) |
| Justified asking<br>husband to use<br>condom if he has a<br>sexually transmitted<br>disease | No / Does not know/ Missing | 18.2% (2.3%-60.6%) | 11.5% (1.5%-29.8%) |
|  | Yes | 81.8% (39.4%-97.7%) | 88.5% (70.2%-98.5%) |
| Ever tested for HIV | No/ Missing | 50.8% (13.8%-86.7%) | 68.8% (19.1%-92.6%) |
|  | Yes | 49.2% (13.3%-86.2%) | 31.2% (7.4%-80.9%) |

The median, minimum and maximum of percentages among all countries are shown.

We found both differences and similarities in crude odds ratios between variables across the sexes. Figure 1 shows the associations between all pairs of variables, each rectangle showing the crude odds ratios for the 29 countries. For example, for both sexes, false beliefs about AIDS were positively associated with rural location and judging gender-based violence justified (represented by red rectangles in the figure). Additionally, for women false beliefs about AIDS were positively associated with sexual initiation before age 16. Having ever been tested for HIV was positively associated for both sexes with the belief that it is justified for a wife to ask the husband to use a condom if he has a sexually transmitted infection (STI), with being married, with media access, and with literacy. These findings were consistent across countries.

**Figure 1.**
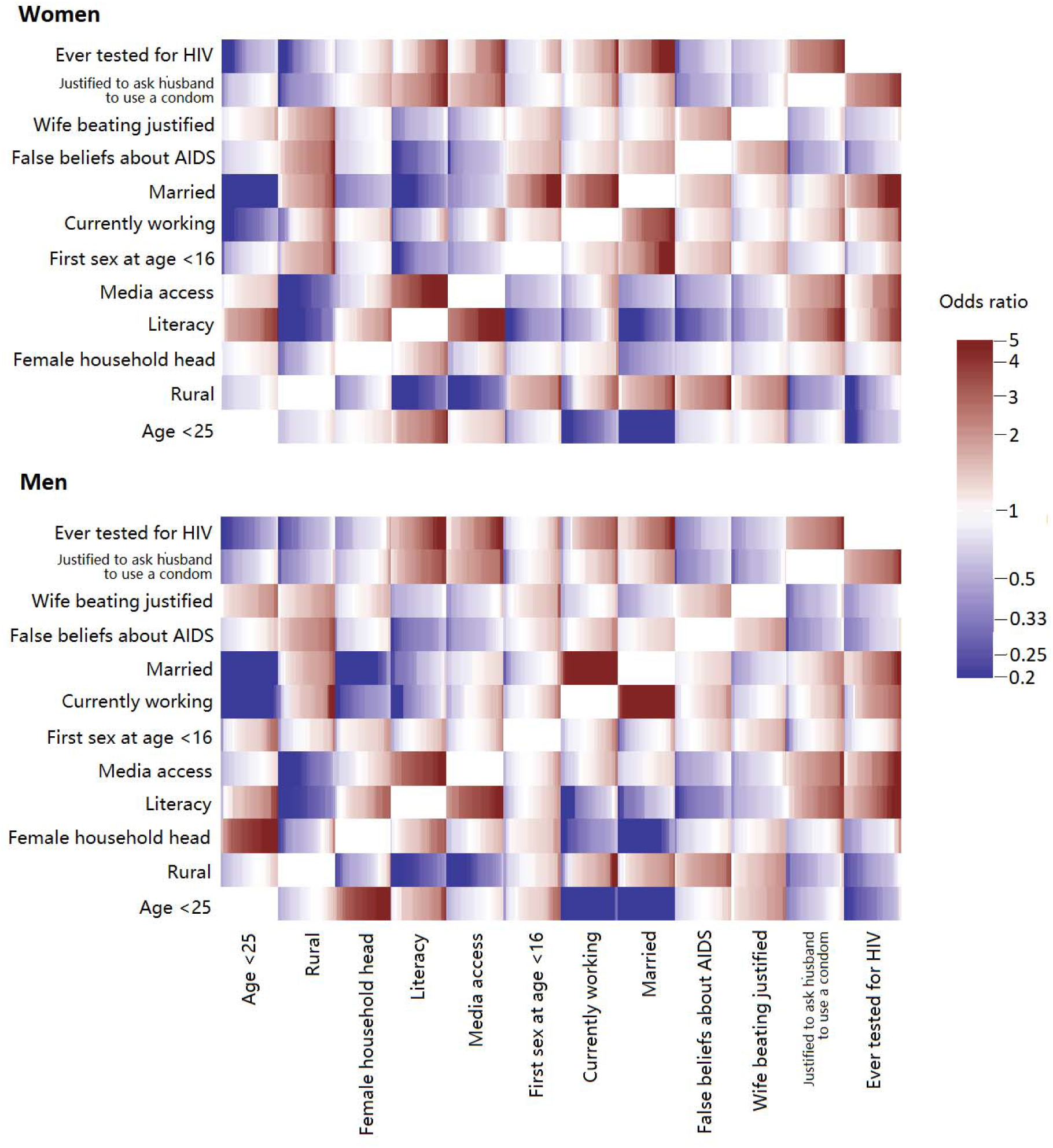
Crude odds ratios for all pairs of selected variables for women (upper panel) and men (lower panel) in the analysis without HIV status (29 countries). Each rectangle shows the odds ratios between the corresponding pair of variables for all countries, sorted from lowest to highest.

Figure 2 shows the estimated summary graphs from the Bayesian network analysis for women and men. The thickness of the edges is proportional to the number of countries for which the given edge was present, and the colour and pattern indicate if the association (i.e. the signs of both crude and conditional log odds ratios) was always negative (blue, dashed), positive (red, solid), or differed between countries (grey, dotted). In both men and women, literacy and urbanity had the largest number of edges in the summary graphs, including many with variables potentially linked with the probability of acquiring HIV. For instance, for women, literacy was negatively associated with sexual initiation before age 16 and false beliefs about AIDS in most countries (21 and 24, respectively). Moreover, for women, literacy was positively associated with the belief that it is justified to ask the husband to use a condom if he has an STI (12 countries) and having ever been tested for HIV (8 countries). For men, literacy was positively associated with having been tested for HIV and negatively associated with having false beliefs about AIDS in almost all countries (20 and 27 countries, respectively). In 13 countries, literacy among men was also positively associated with believing the wife is justified to ask the husband to use a condom if he has an STI. For both men and women, living in a rural area was positively associated with accepting domestic violence against the wife (women 16 countries, men 9 countries) and having false beliefs about AIDS (14 countries for both men and women), and negatively associated with having been tested for HIV (women 12 countries, men 11 countries). For women, the association between rural residence and false beliefs about AIDS was however not present in any country with high (>3%) HIV prevalence. Having been tested for HIV was positively associated with the belief that it is justified to ask the husband to use a condom (women 24 countries, men 8 countries). Having ever been tested for HIV was also associated with having access to media in 8 countries for women and 11 countries for men, all of which were countries with low (<3%) HIV prevalence. Among women, having been tested for HIV also was positively associated with being married in 21 countries. We found only very few directed edges, and will not interpret these in a fully causal way. Detailed results for each country are reported in <u>Additional file 1</u>.

**Figure 2.**
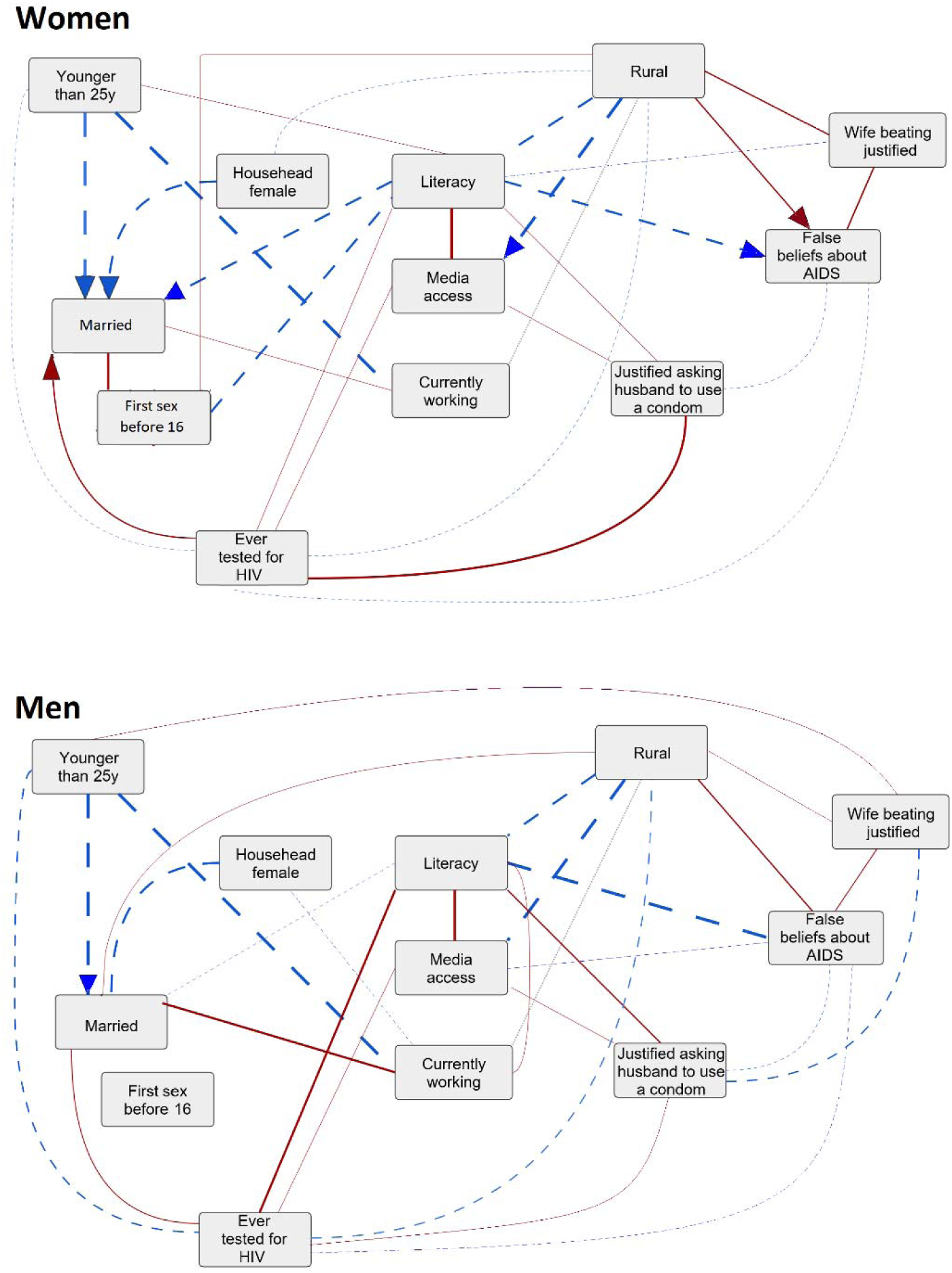
Summary graphs for the Bayesian network analysis without HIV status (29 countries) for women (upper panel) and men (lower panel). Shows associations present in at least six countries. Line thickness is proportional to the number of countries where the edge was present. The edges are oriented (arrowheads) if the number of countries with the shown direction was at least six higher than the number of countries with the opposite direction. Blue dashed lines indicate negative conditional associations, red solid lines indicate positive conditional associations and grey dotted lines indicate that the sign of the association differed across countries.

HIV data were unavailable for Benin, Congo, Kenya, Mozambique, Nigeria and Uganda. In the analysis of the remaining 23 countries where we included HIV status, testing HIV positive was negatively associated with younger age and living in a rural location for both sexes (Figure 3). Being HIV infected was positively associated with being married, thinking it is justified for a wife to ask a husband to use a condom if he has an STI, and having ever been tested for HIV for both sexes. For women, having a female household head was also positively associated with HIV positivity.

**Figure 3.**
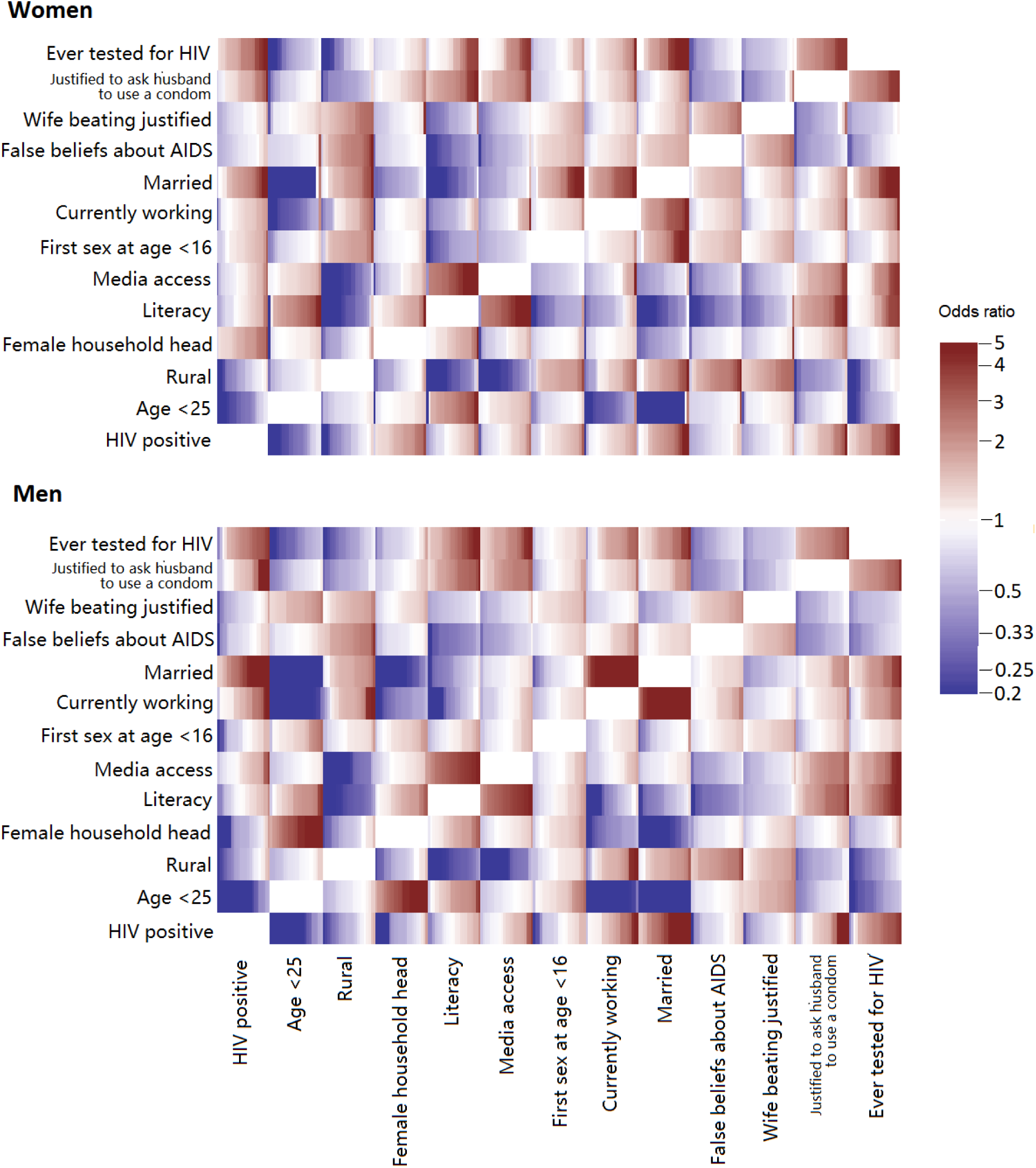
Crude odds ratios for all pairs of the selected variables for women (upper panel) and men (lower panel) in the analysis with HIV status (23 countries). Each rectangle shows the odds ratios between the corresponding pair of variables for all countries, sorted from lowest to highest.

For men, adding HIV status yielded little difference: HIV status was not connected with any other variable in the summary graph (Figure 4). For women, being HIV positive was positively associated with having a female household head, and negatively associated with younger age and living in a rural area, in at least five countries.

**Figure 4.**
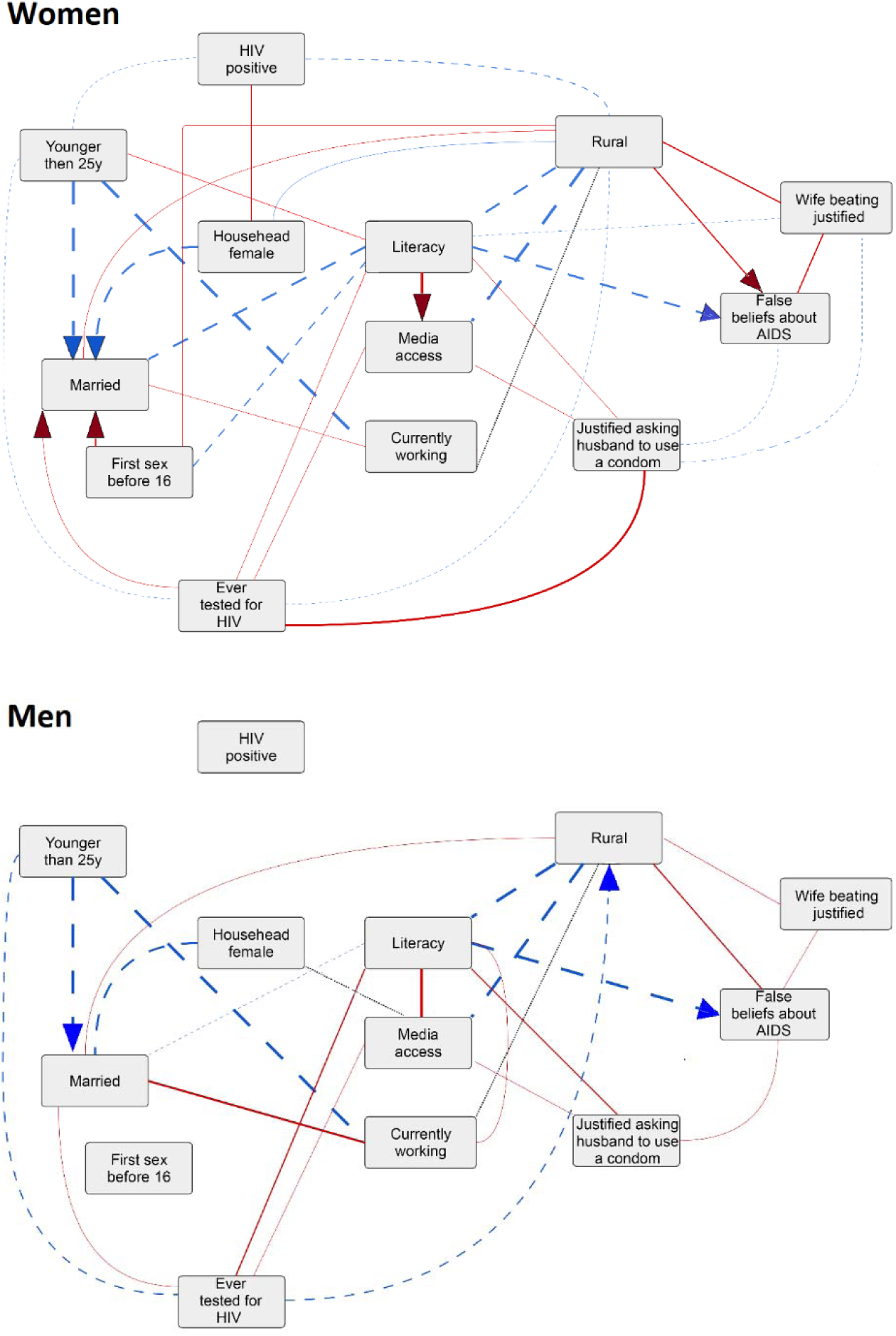
Summary graphs for the Bayesian network analysis with HIV status (23 countries) for women (upper panel) and men (lower panel). Shows associations present in at least five countries. Line thickness is proportional to the number of countries where the edge was present. The edges are oriented (arrowheads) if the number of countries with the shown direction was at least five higher than the number of countries with the opposite direction. Blue dashed lines indicate negative conditional associations, red solid lines indicate positive conditional associations and grey dotted lines indicate that the sign of the association differed across countries.

## Discussion

Our analysis revealed a detailed picture of crude and fully adjusted associations between many socio-behavioural variables that may be linked to HIV in sub-Saharan African countries. Literacy and urbanity were the most central variables: they were directly connected to at least half of the remaining variables for both men and women in at least six countries. Access to media, information on HIV and HIV testing differed greatly between urban and rural locations. Having false beliefs about AIDS was strongly connected with rural location, being illiterate and accepting domestic violence against women. The likelihood of ever being tested for HIV was higher among people who were married, literate, or living in urban settings. For women, HIV testing was also positively associated with feeling justified to ask the husband to use a condom if he had an STI. Many associations were also observed in only few countries. Only very few directed edges were found.

Literacy was directly associated with several factors that should protect against acquiring HIV. For example, literate women started sexual relationships later, and literate people were less likely to have false beliefs about AIDS. This is promising, since literacy is one of the few variables in our analysis that can be directly improved by interventions. The overall proportion of literacy in sub-Saharan Africa has been slowly increasing over the last decades, from 63% in men and 42% in women in 1990 to 72% in men and 57% in women in 2016 [27]. The literacy rates among young men and women are considerably higher and likely to continue to rise.

Earlier studies have returned conflicting results about the positive association between literacy and HIV. For example, a study from Mozambique showed that literacy was associated with HIV knowledge among women living in rural settings [28]. In the Gambia, both the degree of literacy and level of education were positively associated with adherence to therapy among patients receiving ART [29]. In our study, illiteracy was also strongly associated with false beliefs about AIDS; both are likely linked to a low level of knowledge about HIV in general. For example, in Malawi, people who knew more about HIV prevention were more likely to reject, rather than endorse, myths and misconceptions about HIV transmission [30,31]. But some studies suggest the opposite. A recent review found that highly educated people were at higher risk of HIV. This could stem from a variety of causes, since literacy often correlates with other factors like wealth, type of employment and travel, which may be associated with riskier sexual behaviour. Literacy is also strongly associated with living in an urban setting. Literate people may be more likely to access HIV care and treatment, prolonging survival and thus raising prevalence. Finally, the distribution of HIV burden may depend on the overall wealth and situation in the country. One study showed that in poorer sub-Saharan African countries, wealthier individuals were more likely to be HIV positive, but in wealthier countries, poorer individuals were more likely to be HIV positive [32]. A study from Zimbabwe that used DHS data found that people who could read parts of a sentence had a higher risk of HIV infection than those who could not read at all, or who could read complete sentences [33].

We were slightly surprised by our results on the association between urbanity and variables with protective effects against HIV. HIV prevalence is generally higher in urban than rural settings [34]: for example, in the Democratic Republic of the Congo, distance to a city was negatively associated with HIV [35]. There are several possible explanations for these contradictory findings. Urban residents are likely to be more connected to other people, which may increase the risk of infection, despite other protective factors. HIV may also affect sub-populations differently in different settings; for example, in Kenya the poorest people are more likely to have HIV in urban than rural settings [36]. In Uganda, rural residence was associated with an increase in risky sexual behaviour in people who were on long-term ART [37]. Interestingly, the associations of rural location with less HIV testing and more false beliefs about AIDS were not found in women in high HIV prevalence countries. This may reflect the major role of antenatal care (ANC). The vast majority of women attend at least one ANC visit even in rural settings [38], and in settings with generalized HIV epidemic, HIV testing and counselling is part of routine care.

In most countries in sub-Saharan Africa, HIV testing forms the largest gap in the cascade of care. This gap must be closed to meet the “90-90-90” global treatment target and eventually eradicate the epidemic [39]. We identified many associations between HIV testing uptake and other factors. For women, the strongest associations were with being married and with believing it justified for a woman to ask her husband with an STI to use a condom. These two variables may also be directly linked: a study from Zambia showed that married women who knew that their husbands were either sexually promiscuous or had a sexually transmitted infection were more likely to feel justified in refusing sex or requesting he use condoms [40]. Stigma, fear and discrimination have been identified as barriers for utilizing HIV services. For men, literacy was strongly associated with HIV testing. Although the association may not be causal and men are overall more likely to be literate than women, this shows that men should not be neglected in literacy interventions.

### Strengths and limitations

Our study analysed the associations between socio-behavioural variables in most sub-Saharan African countries. Other epidemiological studies have used Bayesian networks [41–44] but we believe we are the first to investigate detailed associations between variables related to the spread of HIV in sub-Saharan Africa. Our study was also strengthened by our use of a very large data set from the Demographic Health Surveys of almost 30 countries, which allowed us to identify commonalities and differences between countries, and between men and women.

Our study also had several limitations. The variables were pre-selected by our research team, based on the prior knowledge and some practical constraints. Although we assumed there were no hidden variables in our analysis, we had to exclude potentially relevant variables because they were too heterogeneous or data were missing. Religion was associated with certain factors in many countries, but the diversity of religions prevented us from dichotomizing and including this variable. Other variables like male circumcision, ever paying for sex, or using condoms during the last paid intercourse were excluded because they were not collected in some countries or too many values were missing. For example, we could not include HIV status in the main analysis because too little data on HIV-positive men was available. DHS did not collect data on some factors related to geographic variability of HIV and AIDS in sub-Saharan Africa, including sex work, tuberculosis, and use of injected drugs. We had to dichotomize all variables to compute the CPDAGs with the available data, even though for some variables it would have been better to include more categories. For example, for marital status, widowed and divorced were categorized as married. There may also be a risk of differential misclassification in the data: for example, literate individuals with higher education may be more reluctant to admit accepting domestic violence or refusing the use of condom. Finally, we only identified a small number of directed edges. It is possible that most associations were not causal, or it may simply demonstrate how hard it is to prove causal relationships with routine data.

## Conclusions

The Bayesian network approach is a valuable technique for studying the structure of associations between variables from surveys and other data sources. Our findings underline the importance of understanding the role of socio-behavioural and contextual factors in the spread of complex diseases like HIV. The associations with variables related to HIV vary across countries, and it is essential to understand and study further the specific situation in each country to identify possible intervention targets. Literacy and urbanity were associated with several factors that are essential for HIV acquisition. Since literacy is one of the few variables that can be improved by interventions, this seems the most promising intervention target.

## Authors’ contributions

ZB, OK and MM designed the study. ZB and OK reviewed and selected the variables to be included in the analysis. ZB wrote the code and performed the statistical analysis with together with JE, SM, AM and MM. SB, ZB, JE and OK reviewed the relevant literature. ZB and JE wrote a first draft of the manuscript, which was revised by OK. All authors contributed to the interpretation of the results and the final version of the manuscript.

## Acknowledgements

We thank Gilles Kratzer and Reinhard Furrer for helpful discussions and Kali Tal for carefully editing the paper.

## Additional files

Additional file 1: Results of the Bayesian network analysis for each country Text file (pdf).

