## Additional file 1 for "Socio-behavioural characteristics and HIV: findings from a graphical modelling analysis of 29 sub-Saharan African countries"

### Appendix: Results of the Bayesian network analysis for each country

**Legend:**

- (light red) Unoriented edge, crude and conditional odds ratios positive
- (dark red) Oriented edge, crude and conditional odds ratios positive
- (light blue) Unoriented edge, crude and conditional odds ratios negative
- (dark blue) Oriented edge, crude and conditional odds ratios negative

The direction of the oriented edges is from the variable on the vertical axis to the variable on the horizontal axis.

**Table S1. Angola, female**

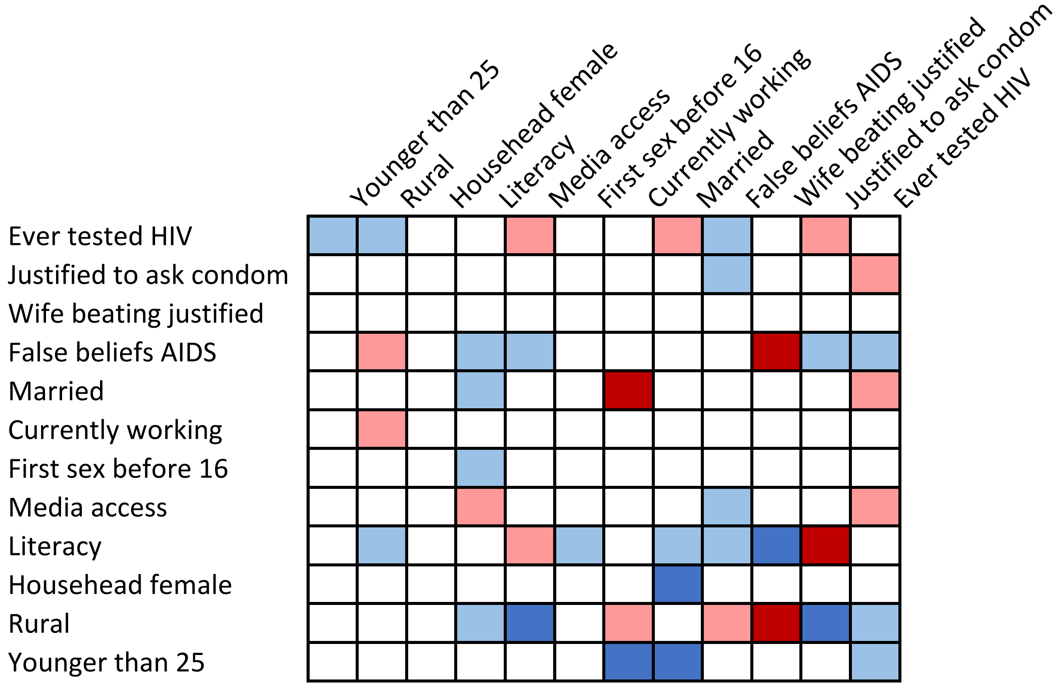


**Table S2. Angola, male**

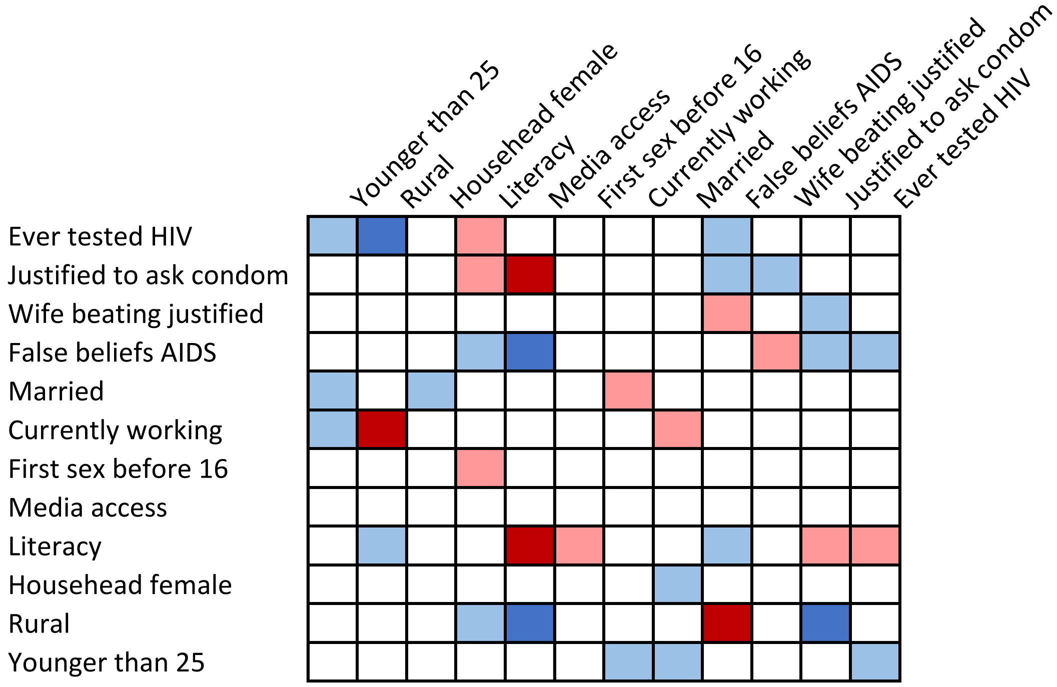

**Table S3. Benin, female**

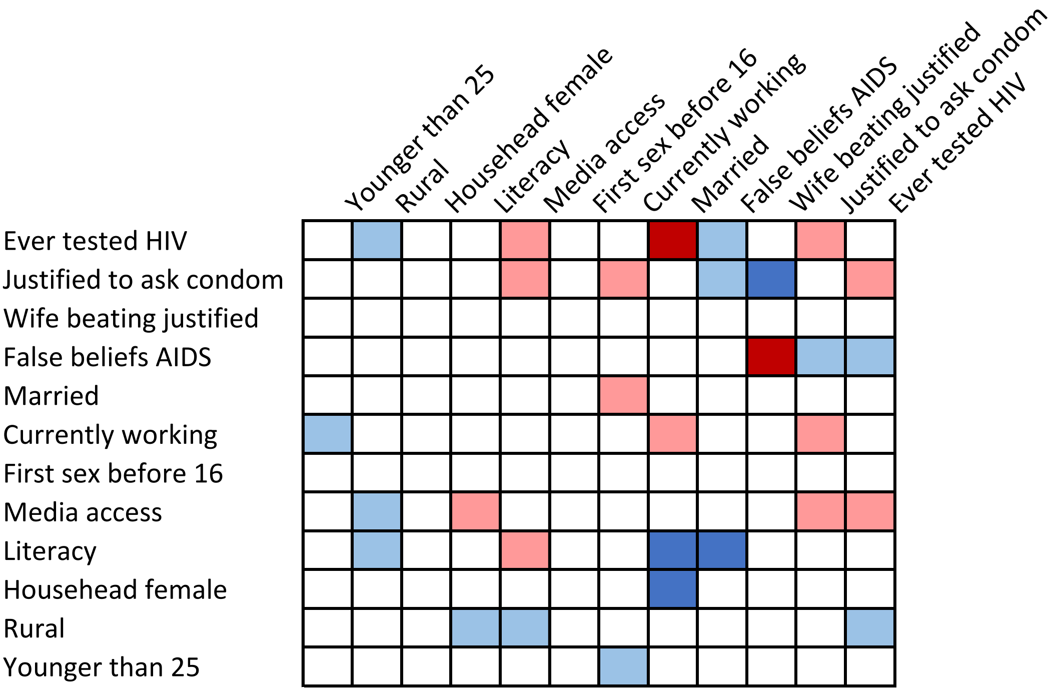


**Table S4. Benin, male**

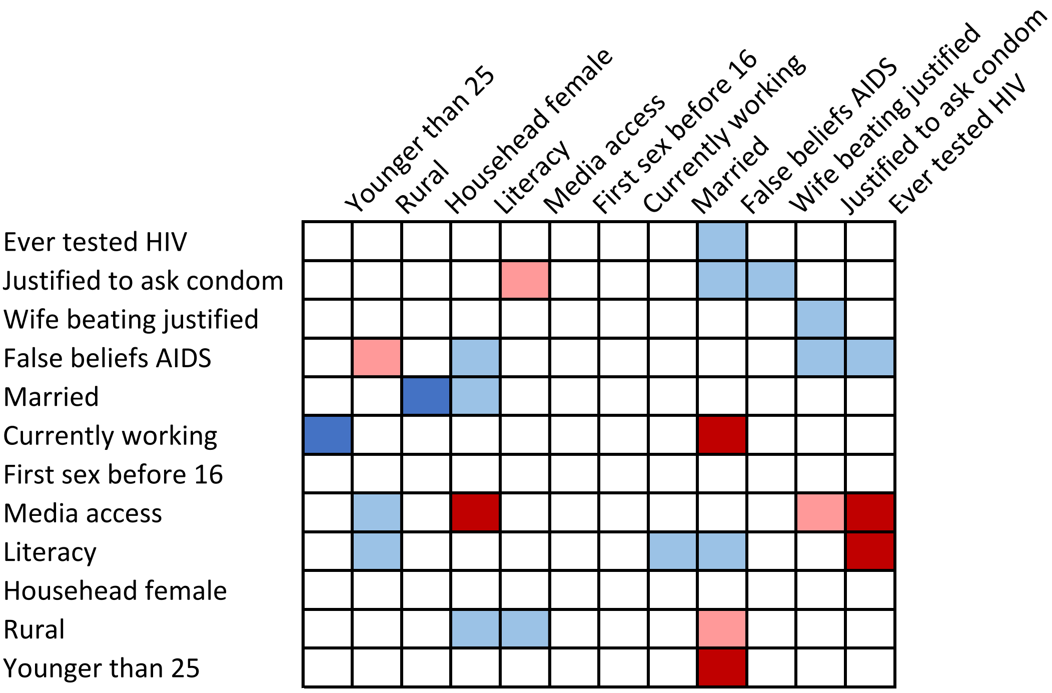

**Table S5. Burkina Faso, female**

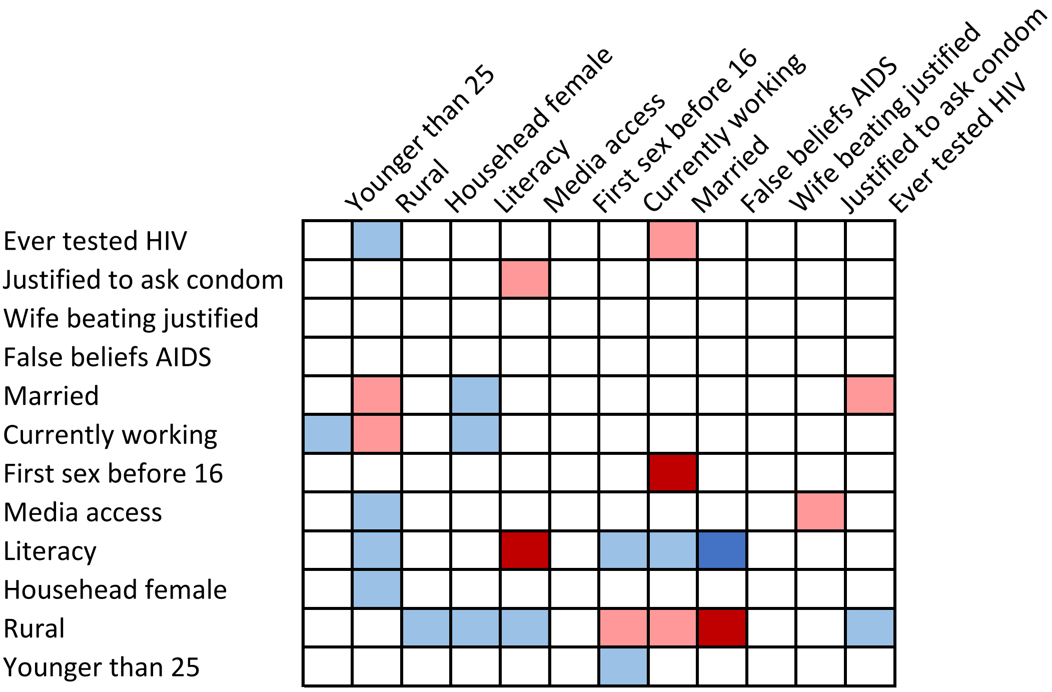


**Table S6. Burkina Faso, male**

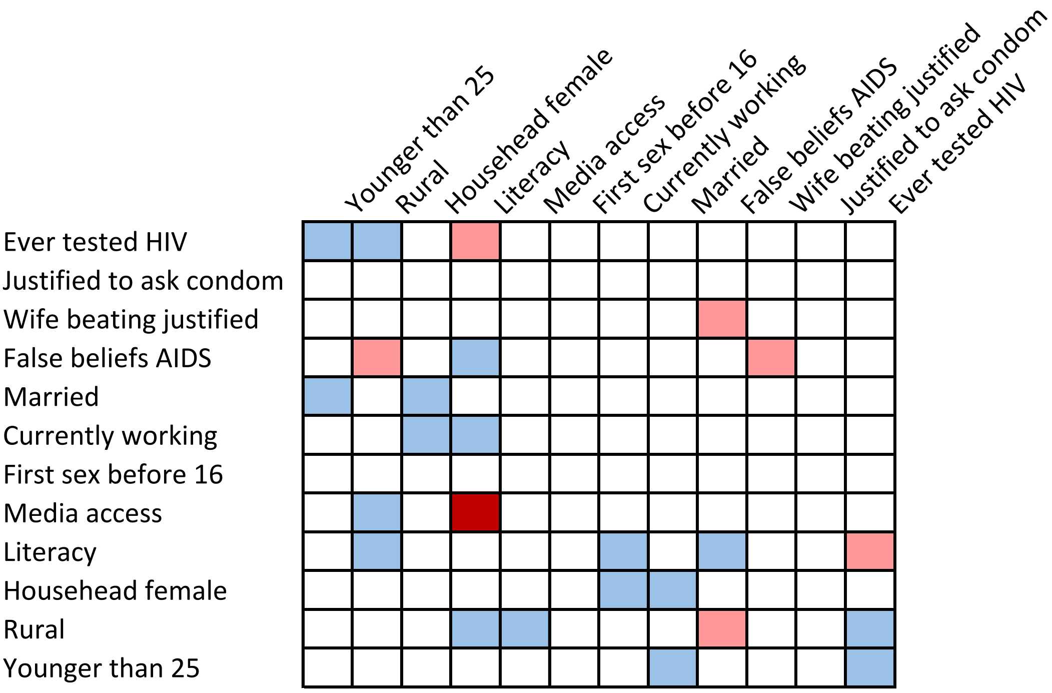

**Table S7. Burundi, female**

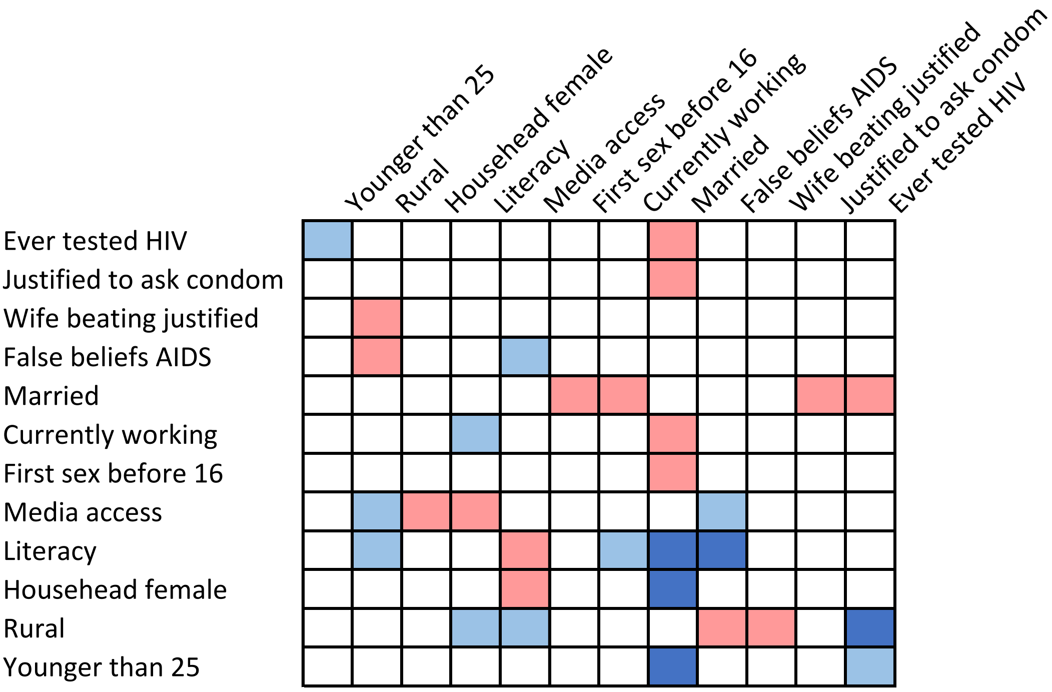


**Table S8. Burundi, male**

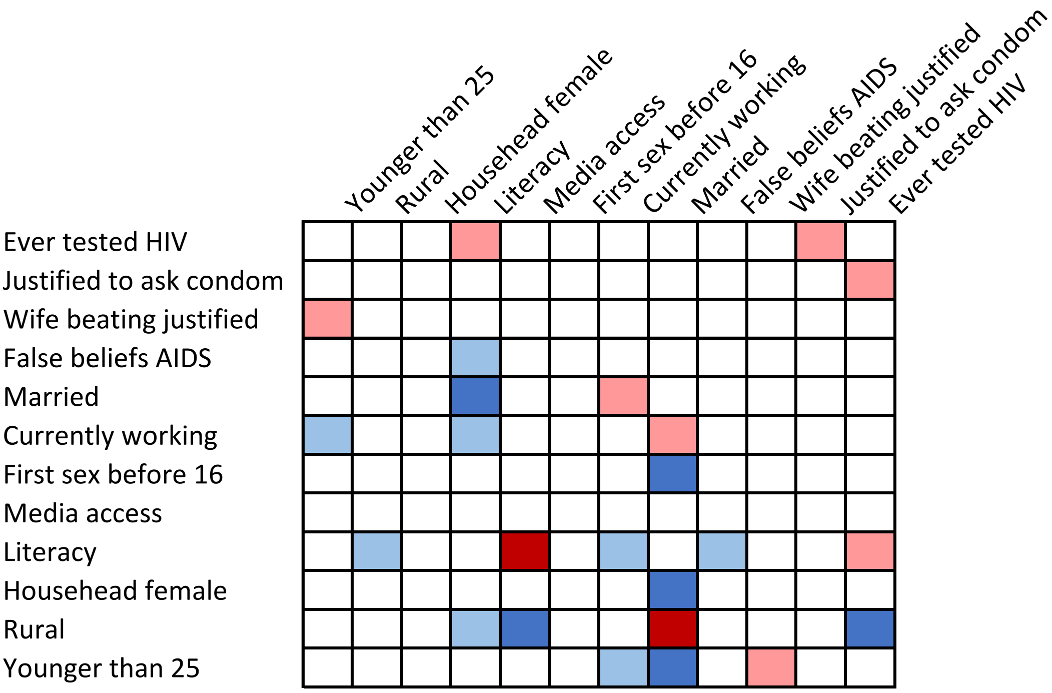

**Table S9. Cameroon, female**

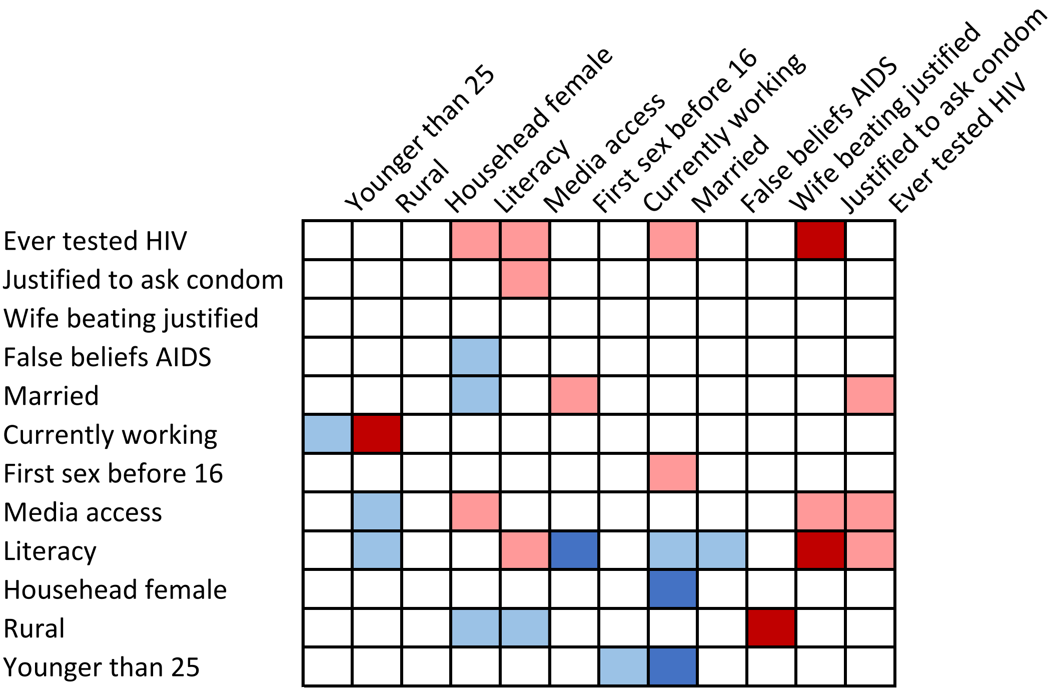


**Table S10. Cameroon, male**

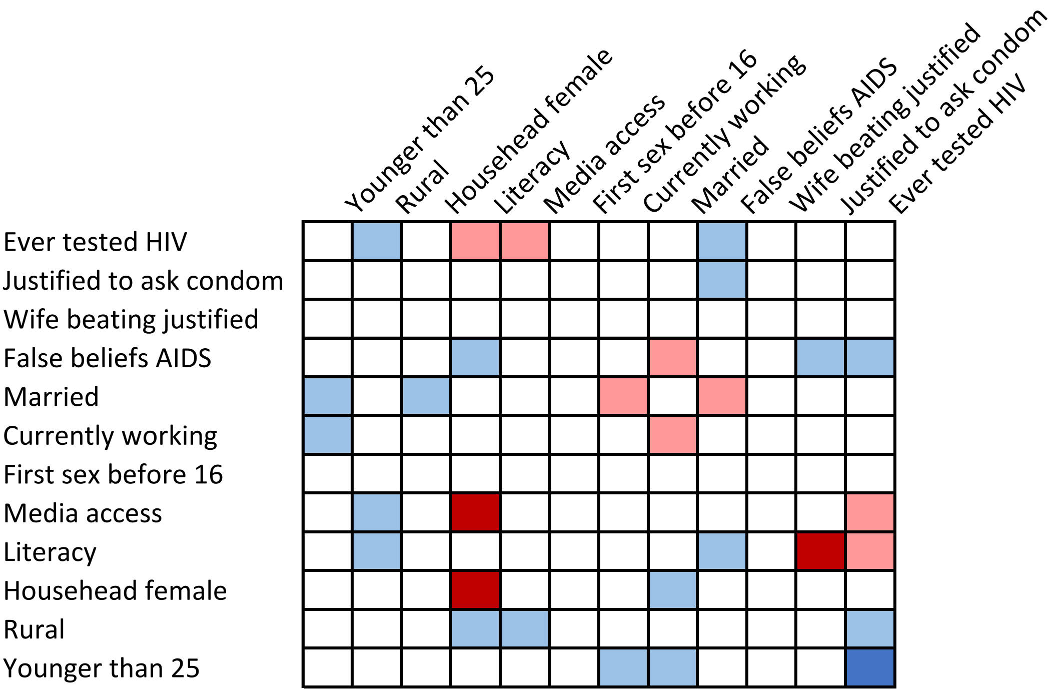

**Table S11. Chad, female**

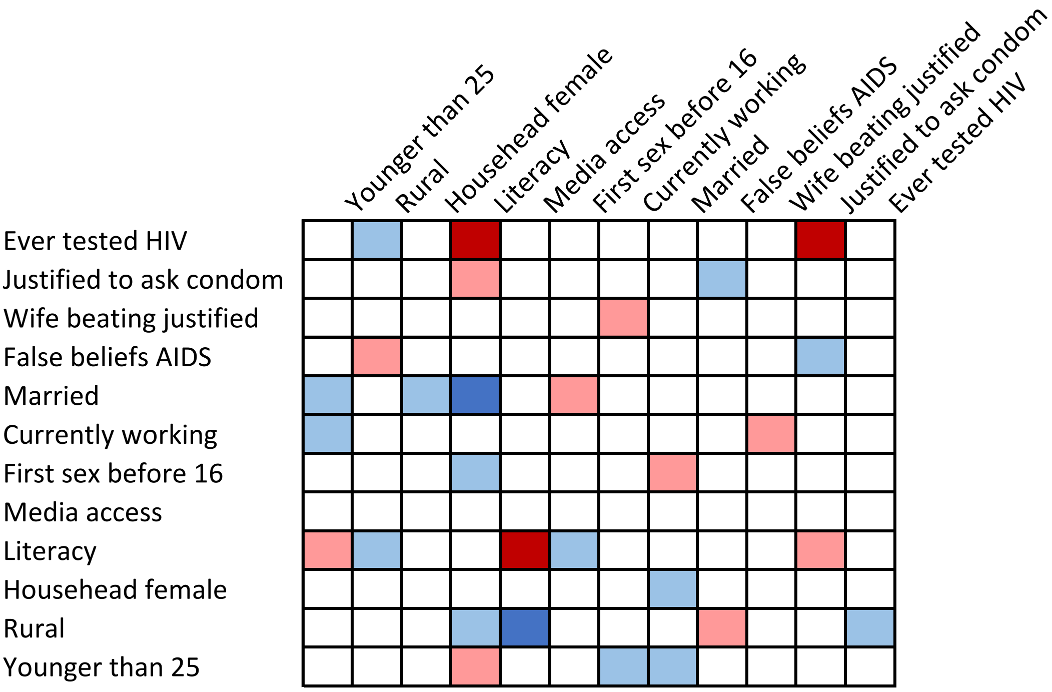


**Table S12. Chad, male**

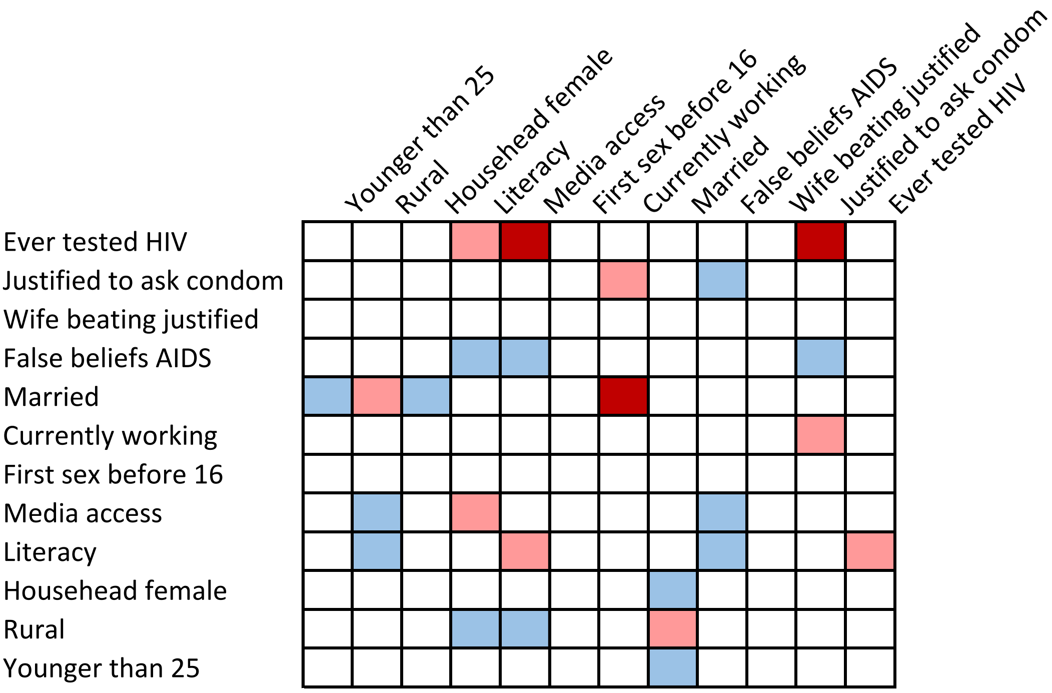

**Table S13. Congo, female**

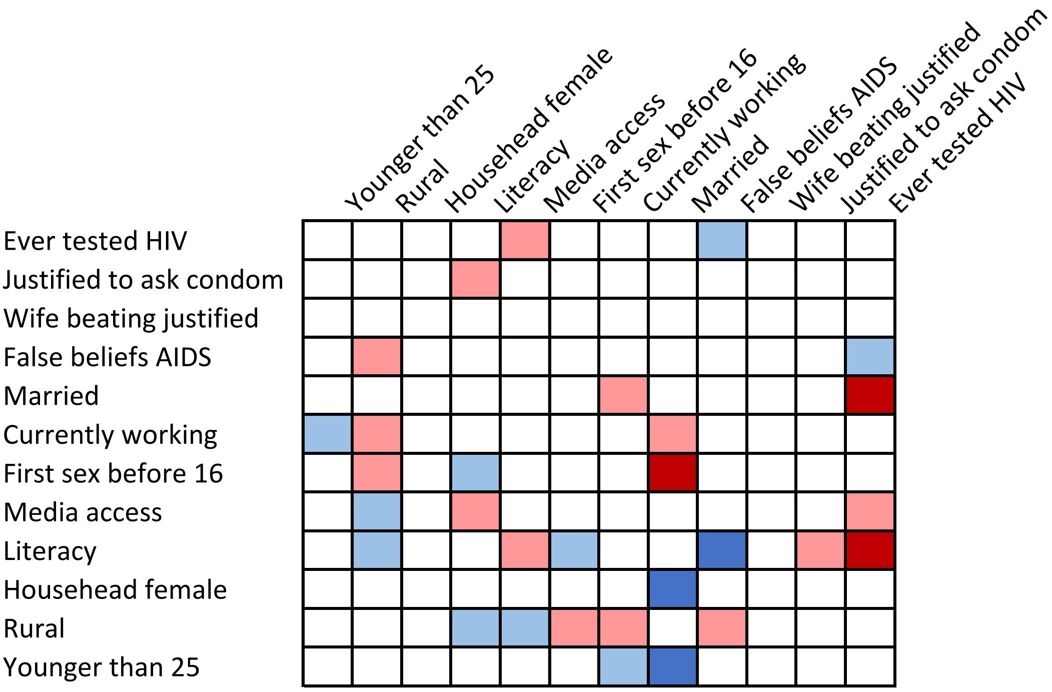


**Table S14. Congo, male**

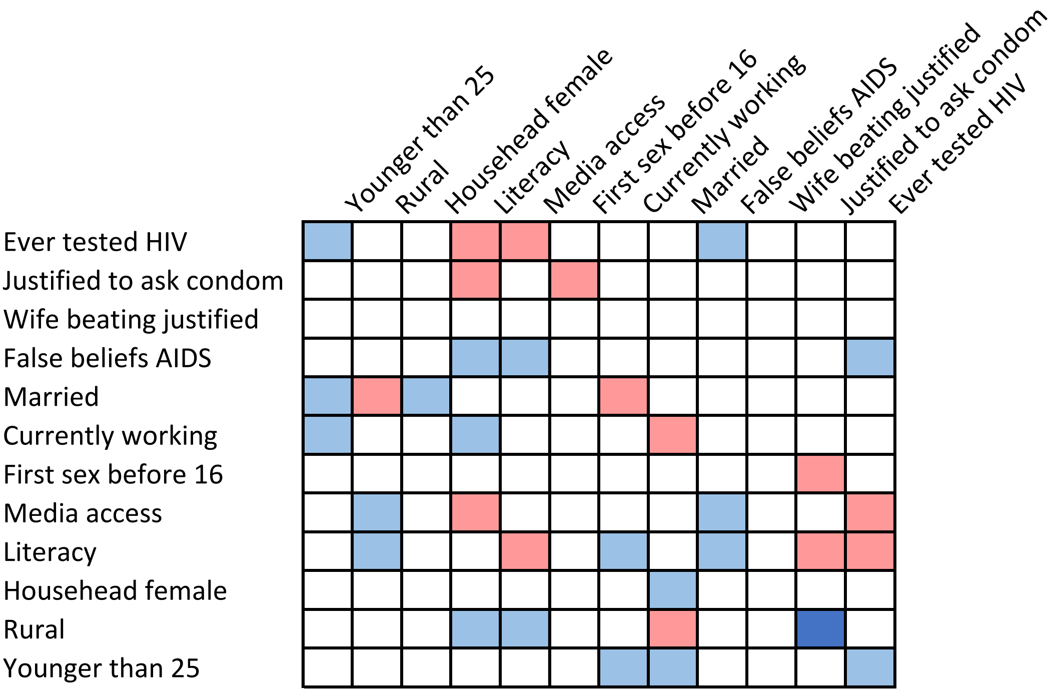

**Table S15. Congo, Democratic Republic of, female**

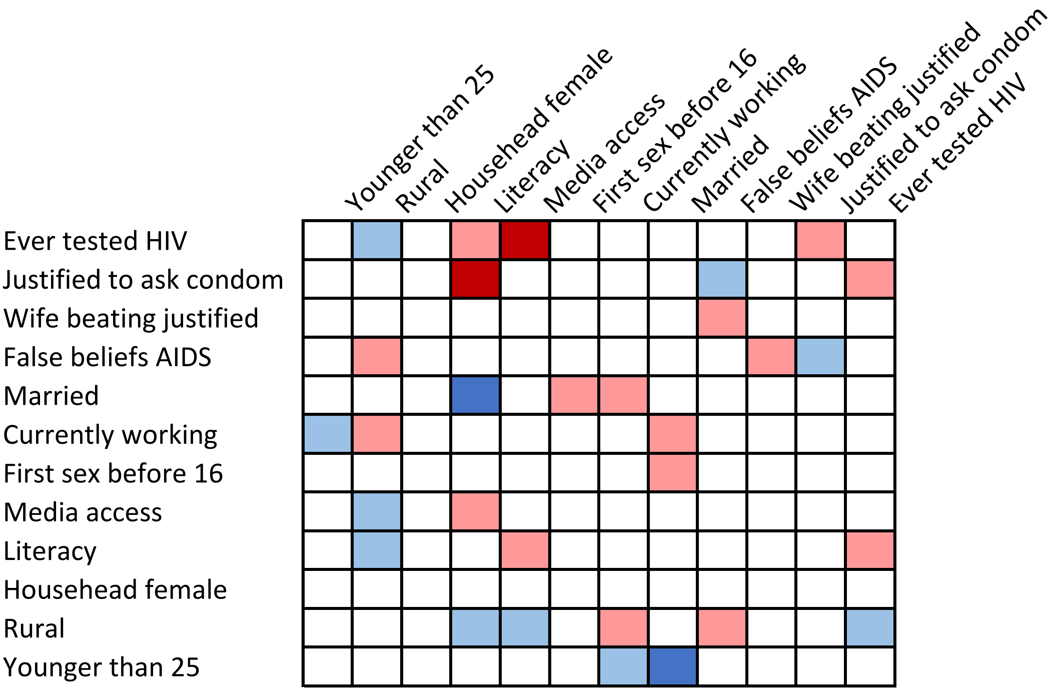


**Table S16. Congo, Democratic Republic of, male**

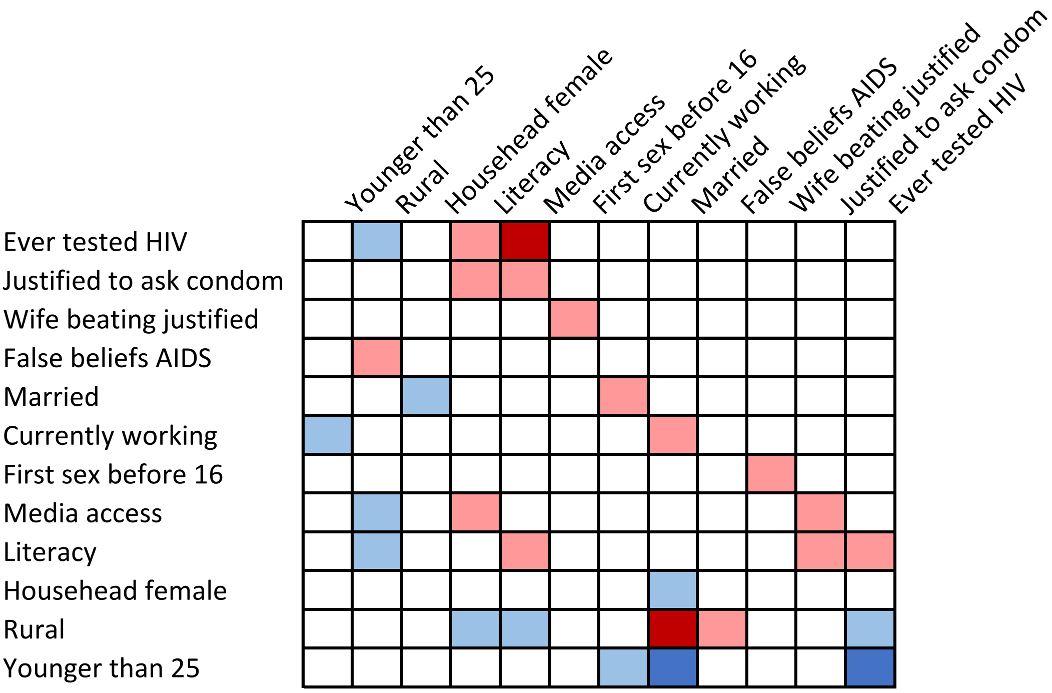

**Table S17. Côte d'Ivoire, female**

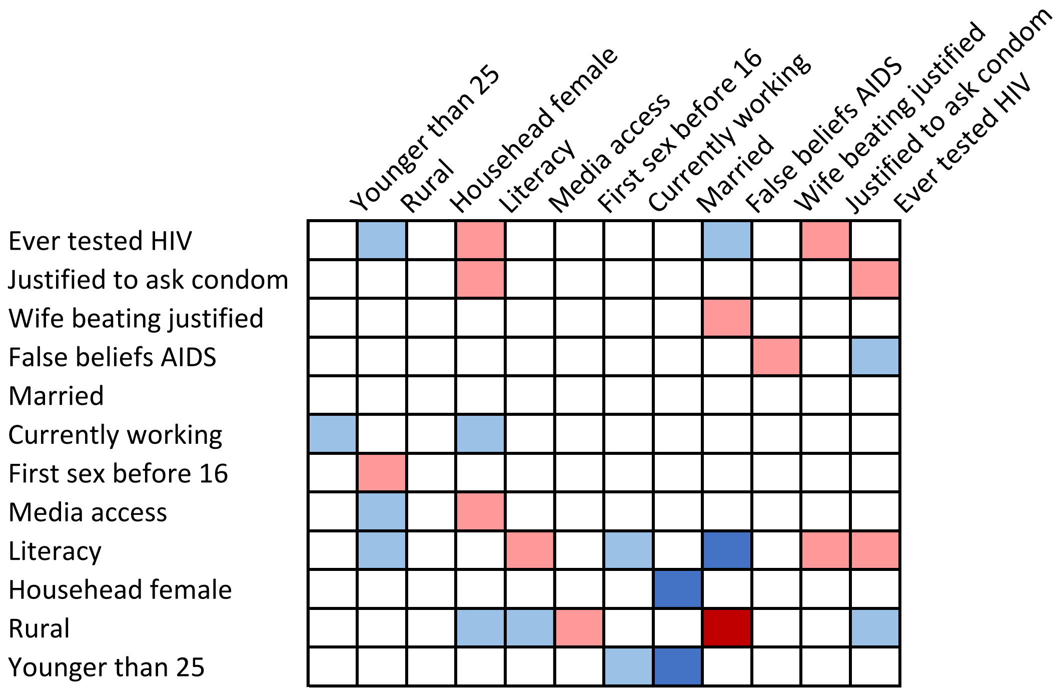


**Table S18. Côte d'Ivoire, male**

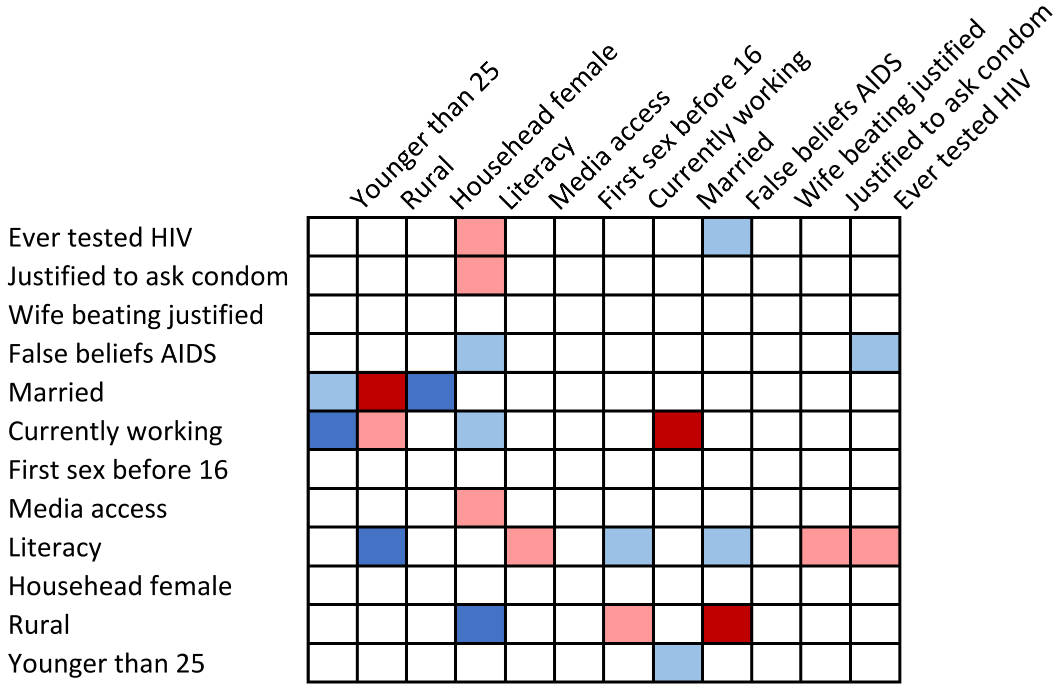

**Table S19. Ethiopia, female**

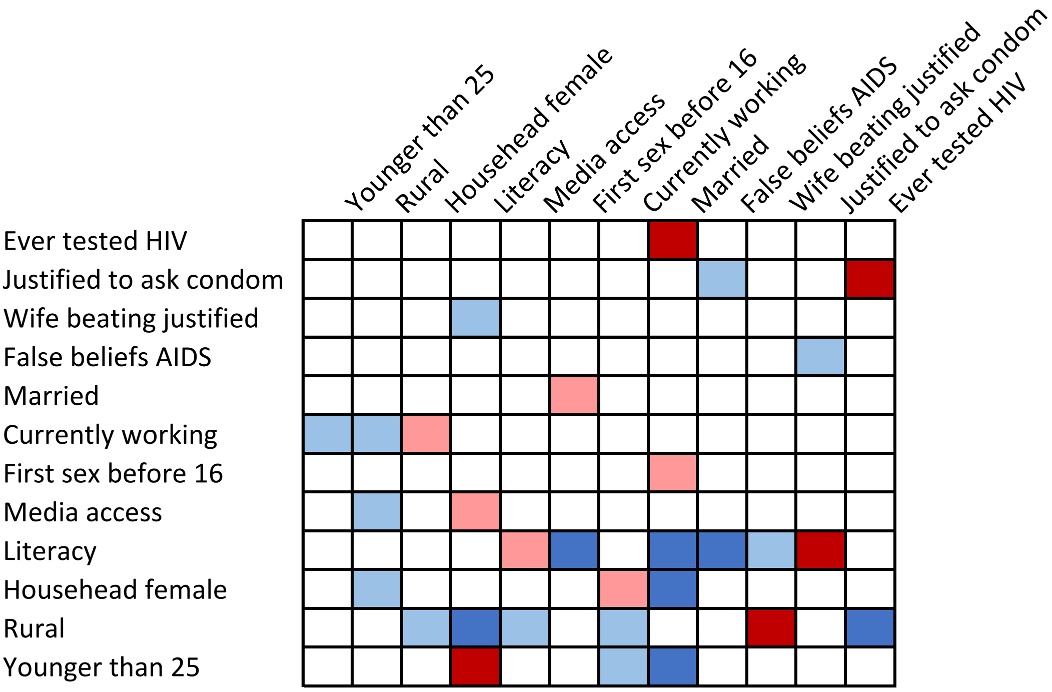


**Table S20. Ethiopia, male**

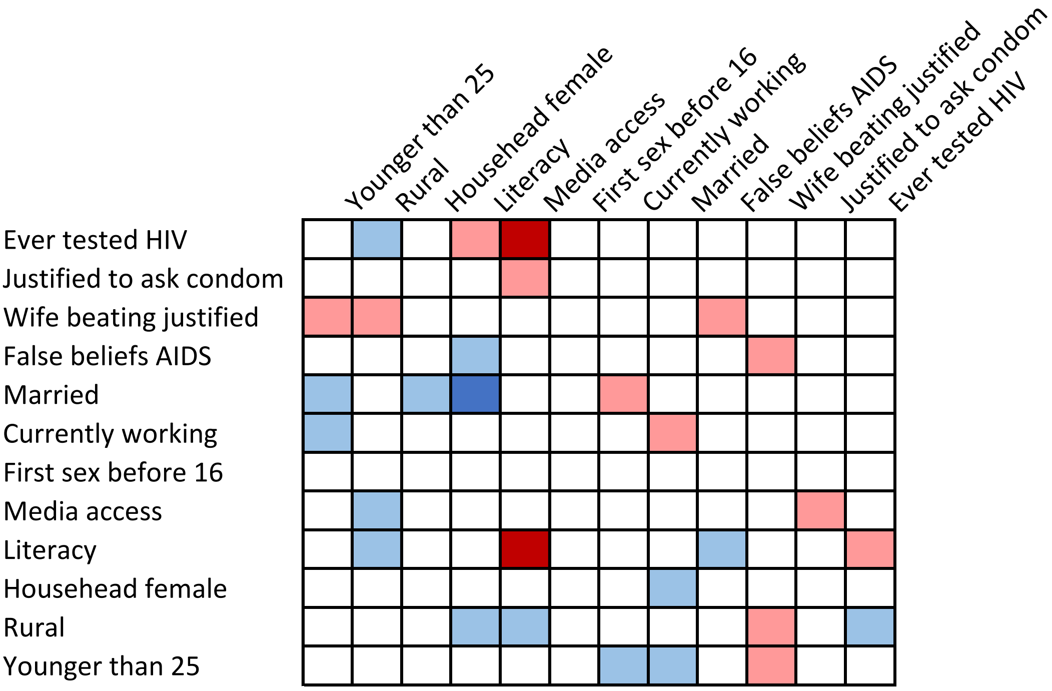

**Table S21. Gabon, female**

| | Younger than 25 | Rural | Househead female | Literacy | Media access | First sex before 16 | Currently working | Married | False beliefs AIDS | Wife beating justified | Justified to ask condom | Ever tested HIV |
|---|---|---|---|---|---|---|---|---|---|---|---|---|
| Ever tested HIV | | | | | | | | | | | | |
| Justified to ask condom | | | | | | | | | | | | |
| Wife beating justified | | | | | | | | | | | | |
| False beliefs AIDS | | | | | | | | | | | | |
| Married | | | | | | | | | | | | |
| Currently working | | | | | | | | | | | | |
| First sex before 16 | | | | | | | | | | | | |
| Media access | | | | | | | | | | | | |
| Literacy | | | | | | | | | | | | |
| Househead female | | | | | | | | | | | | |
| Rural | | | | | | | | | | | | |
| Younger than 25 | | | | | | | | | | | | |

**Table S22. Gabon, male**

| | Younger than 25 | Rural | Househead female | Literacy | Media access | First sex before 16 | Currently working | Married | False beliefs AIDS | Wife beating justified | Justified to ask condom | Ever tested HIV |
|---|---|---|---|---|---|---|---|---|---|---|---|---|
| Ever tested HIV | | | | | | | | | | | | |
| Justified to ask condom | | | | | | | | | | | | |
| Wife beating justified | | | | | | | | | | | | |
| False beliefs AIDS | | | | | | | | | | | | |
| Married | | | | | | | | | | | | |
| Currently working | | | | | | | | | | | | |
| First sex before 16 | | | | | | | | | | | | |
| Media access | | | | | | | | | | | | |
| Literacy | | | | | | | | | | | | |
| Househead female | | | | | | | | | | | | |
| Rural | | | | | | | | | | | | |
| Younger than 25 | | | | | | | | | | | | |

**Table S23. Gambia, female**

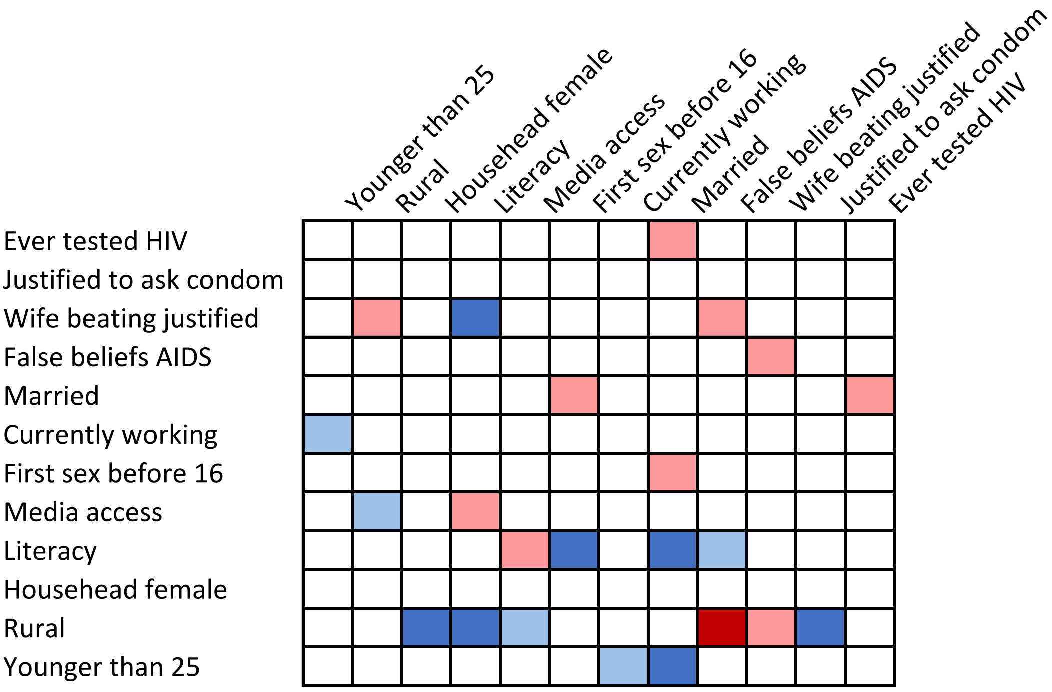


**Table S24. Gambia, male**

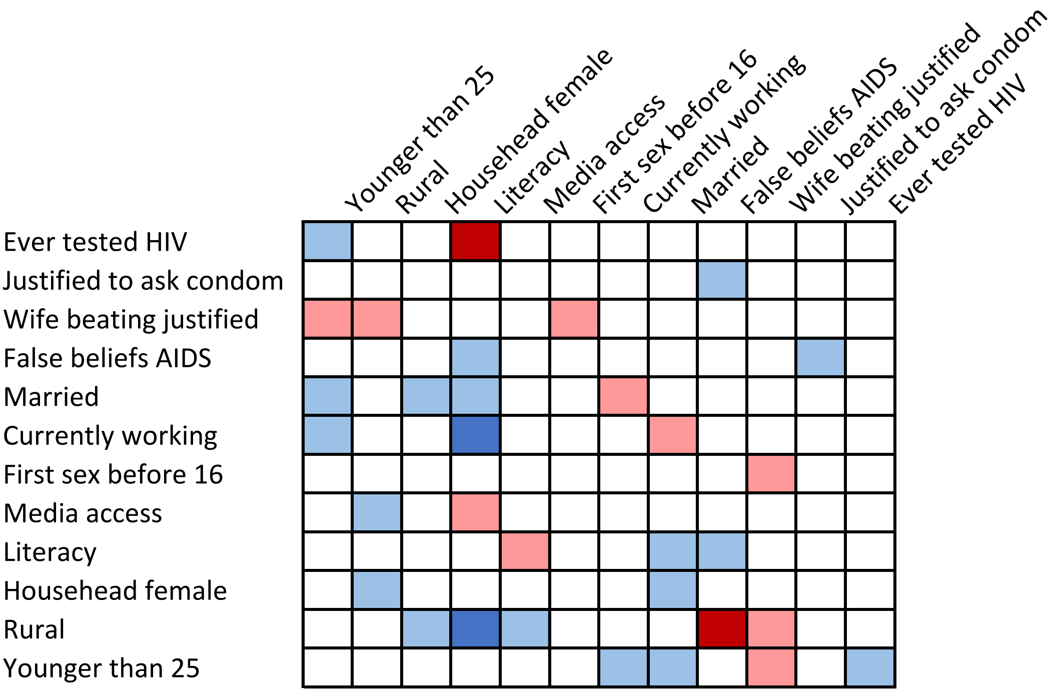

**Table S25. Ghana, female**

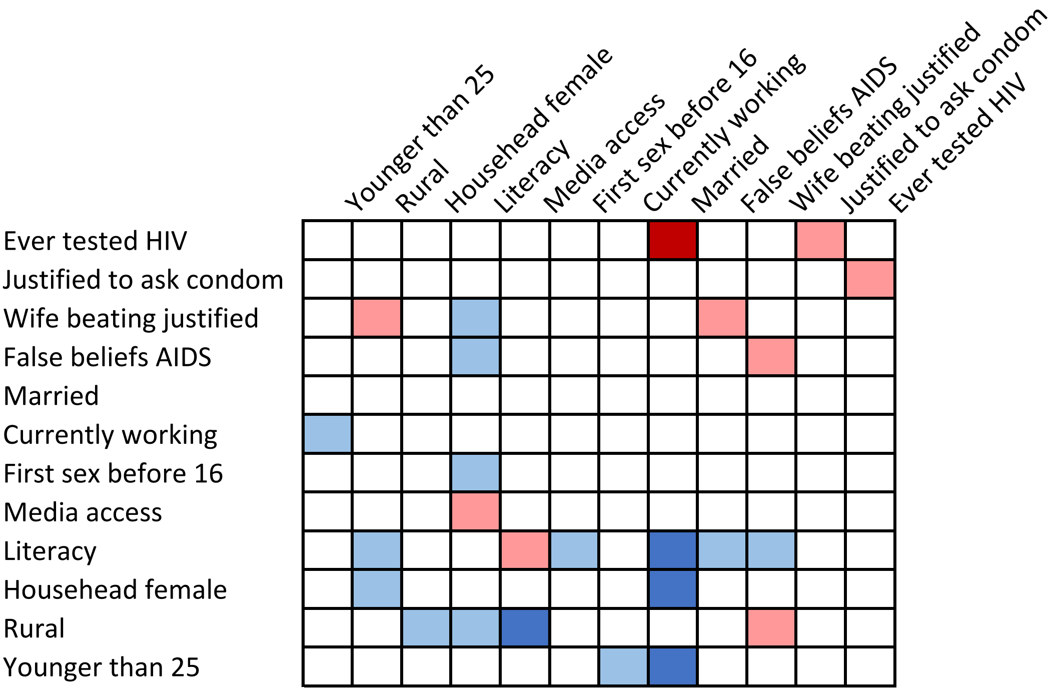


**Table S26. Ghana, male**

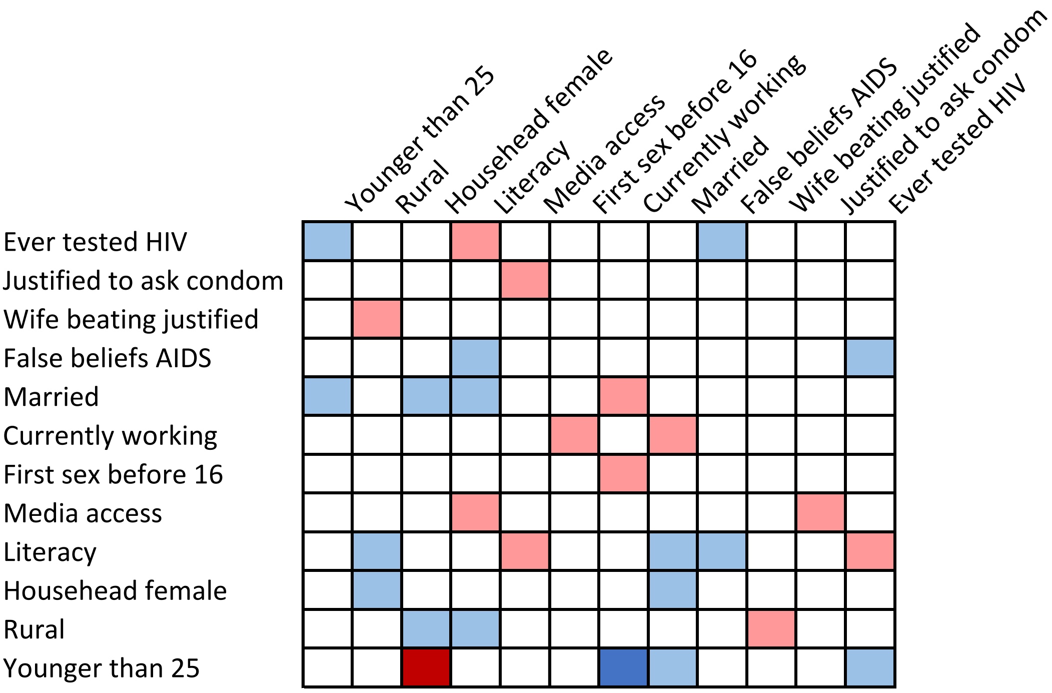

**Table S27. Kenya, female**

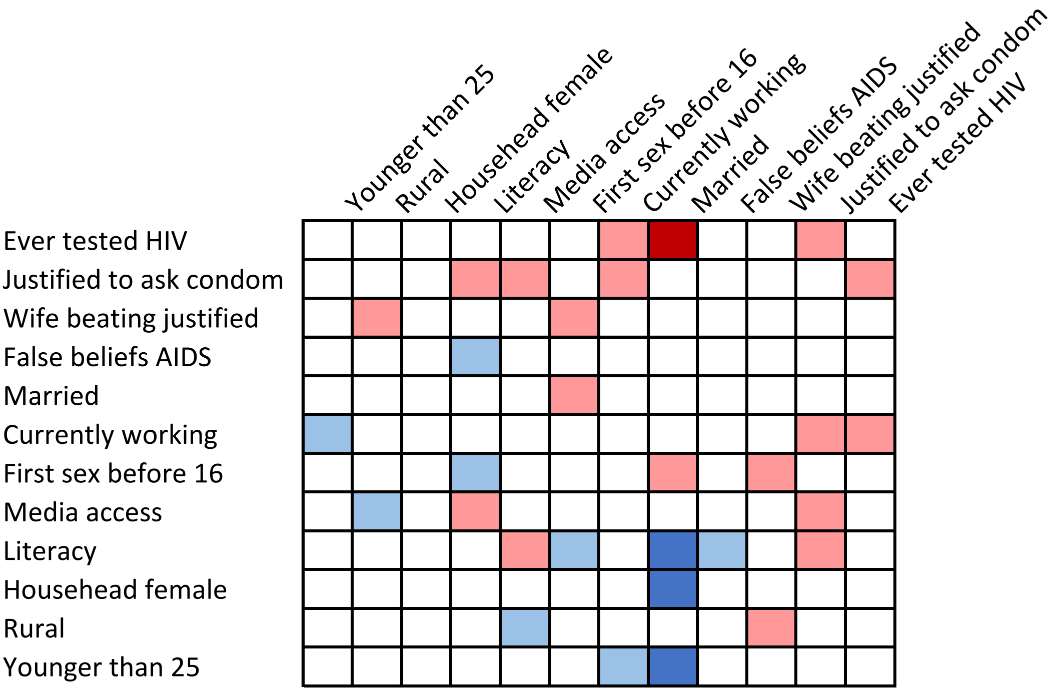


**Table S28. Kenya, male**

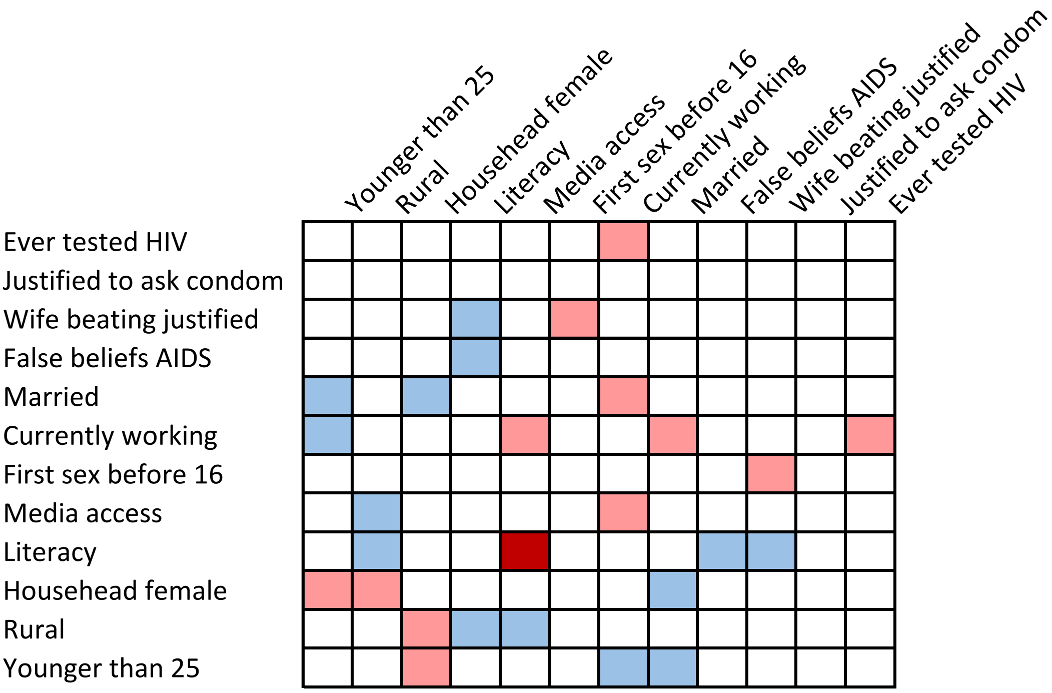

**Table S29. Lesotho, female**

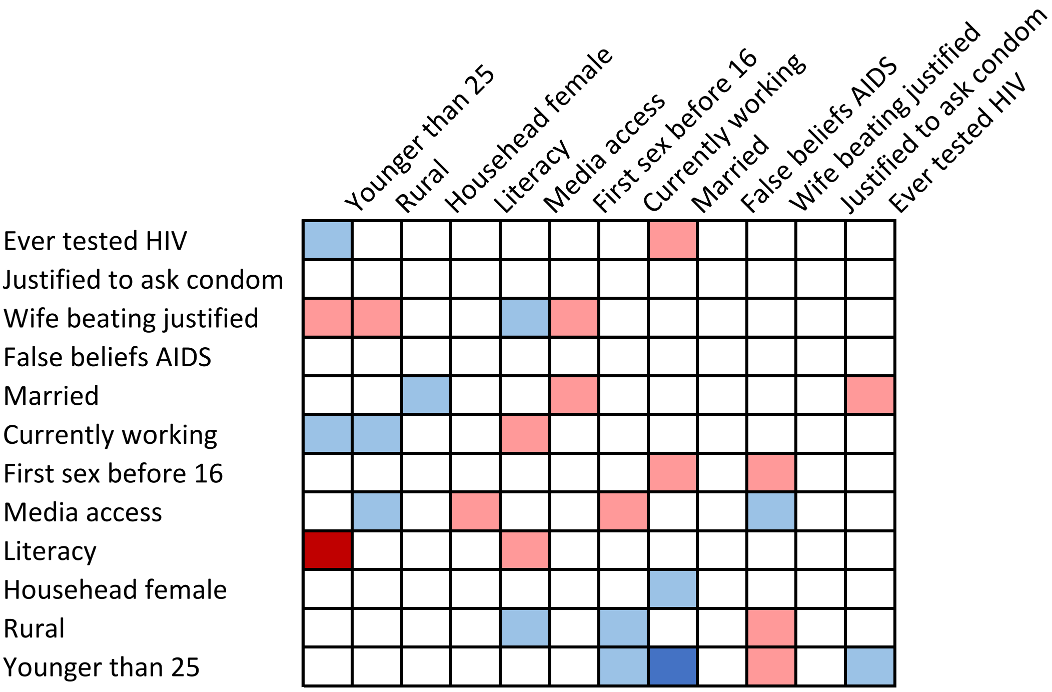


**Table S30. Lesotho, male**

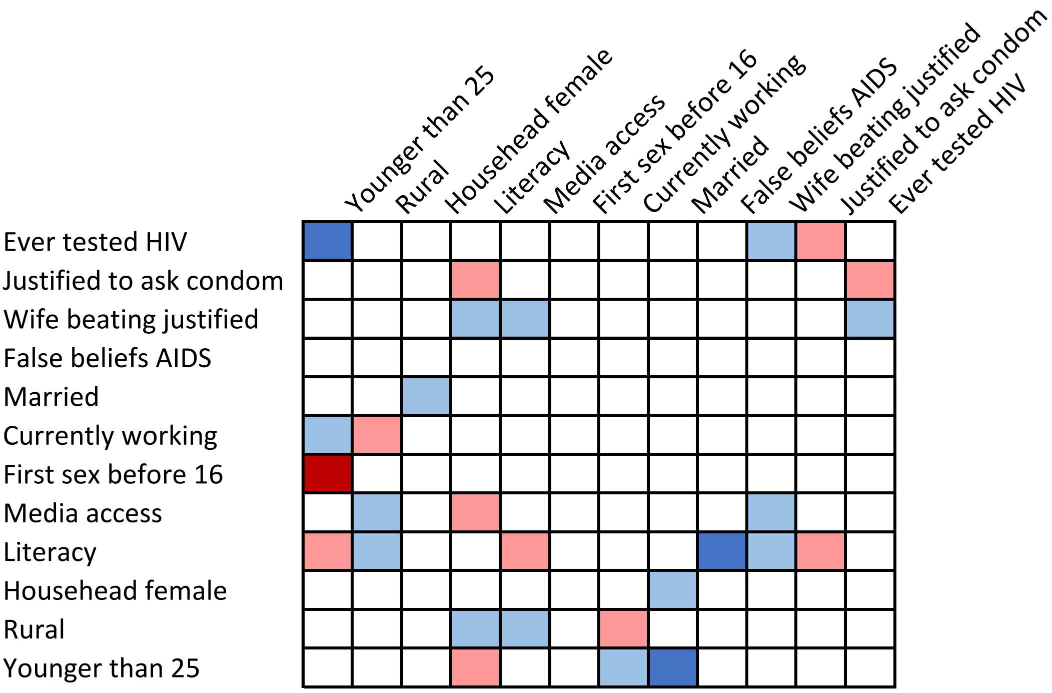

**Table S31. Liberia, female**

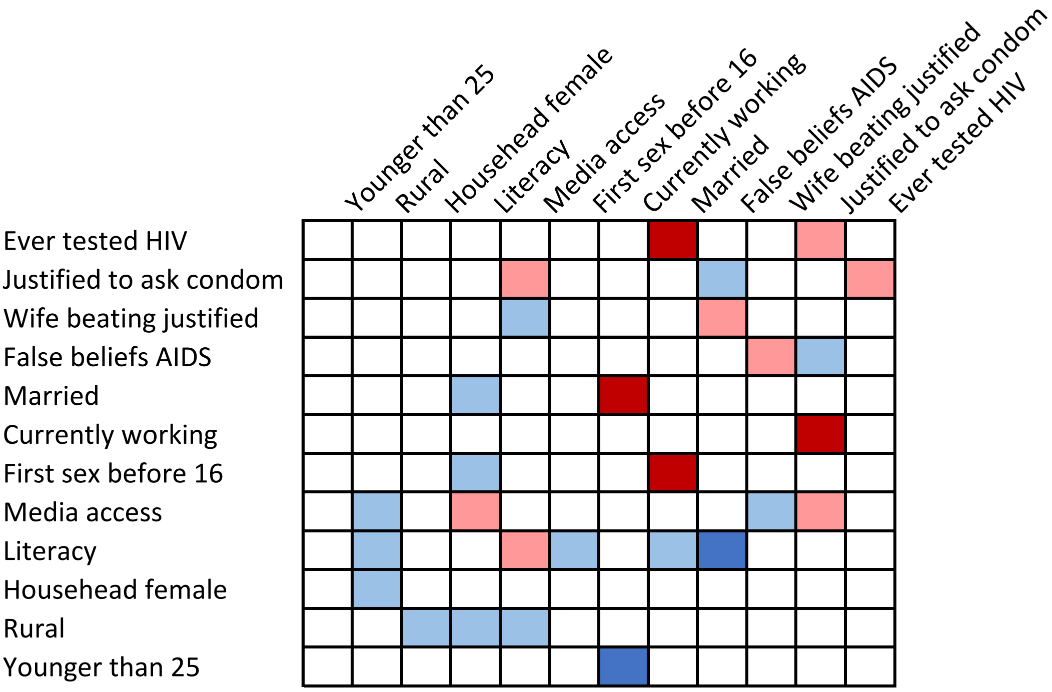


**Table S32. Liberia, male**

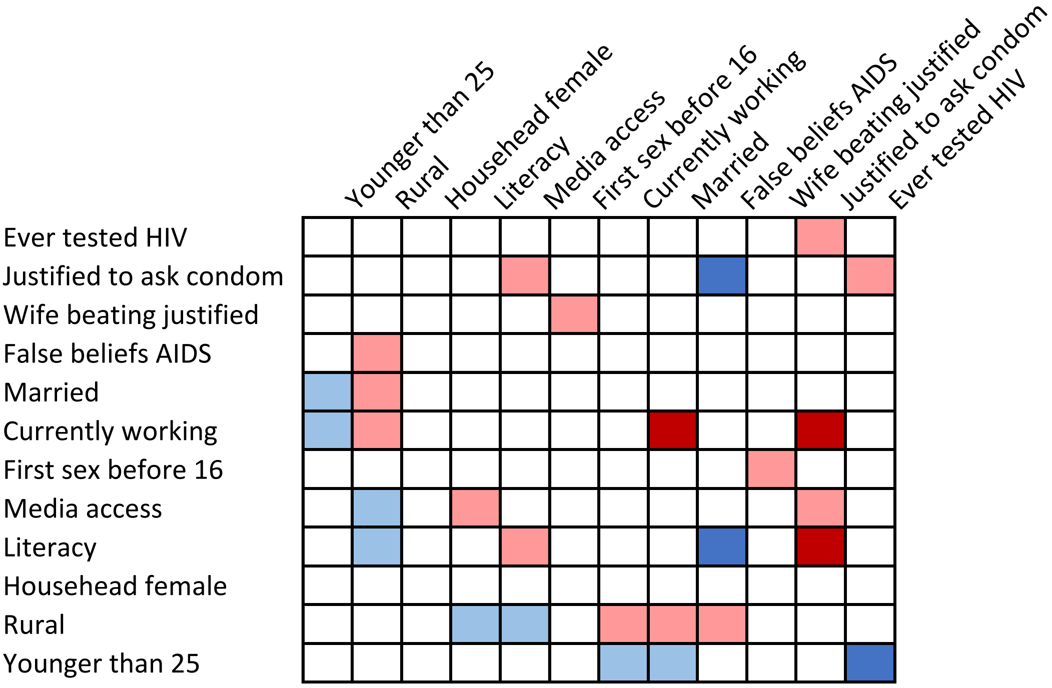

**Table S33. Malawi, female**

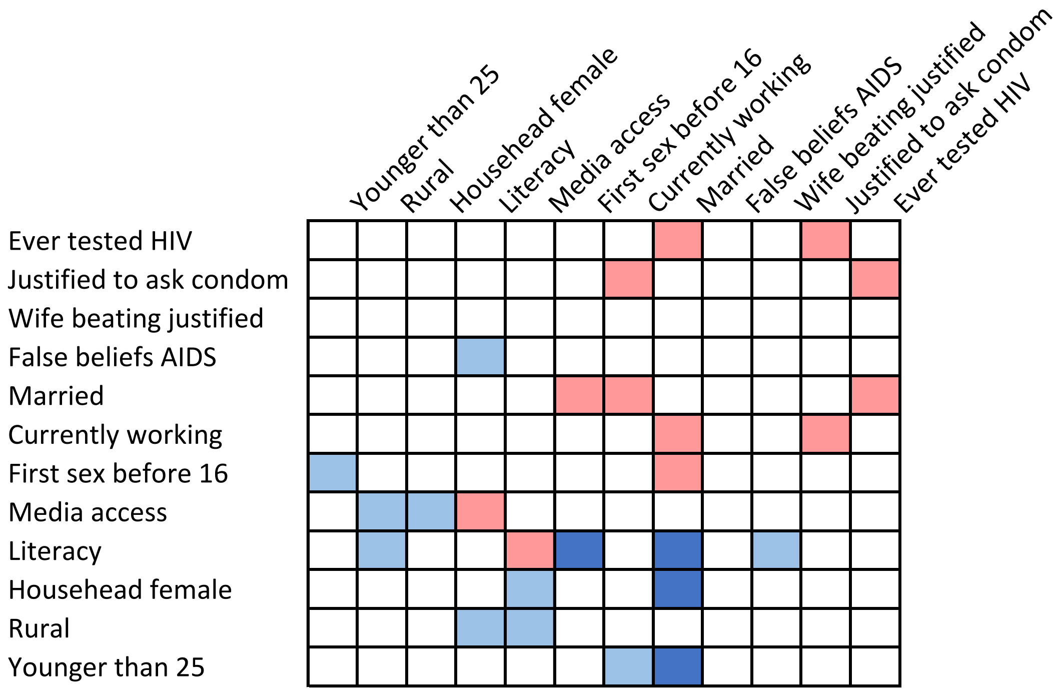


**Table S34. Malawi, male**

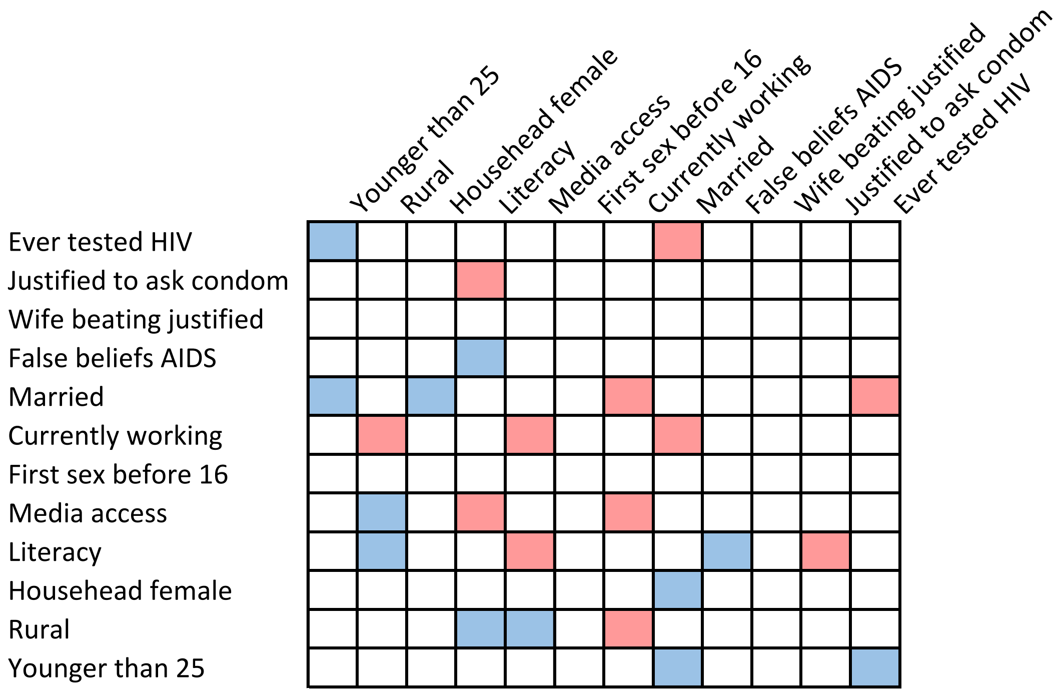

**Table S35. Mali, female**

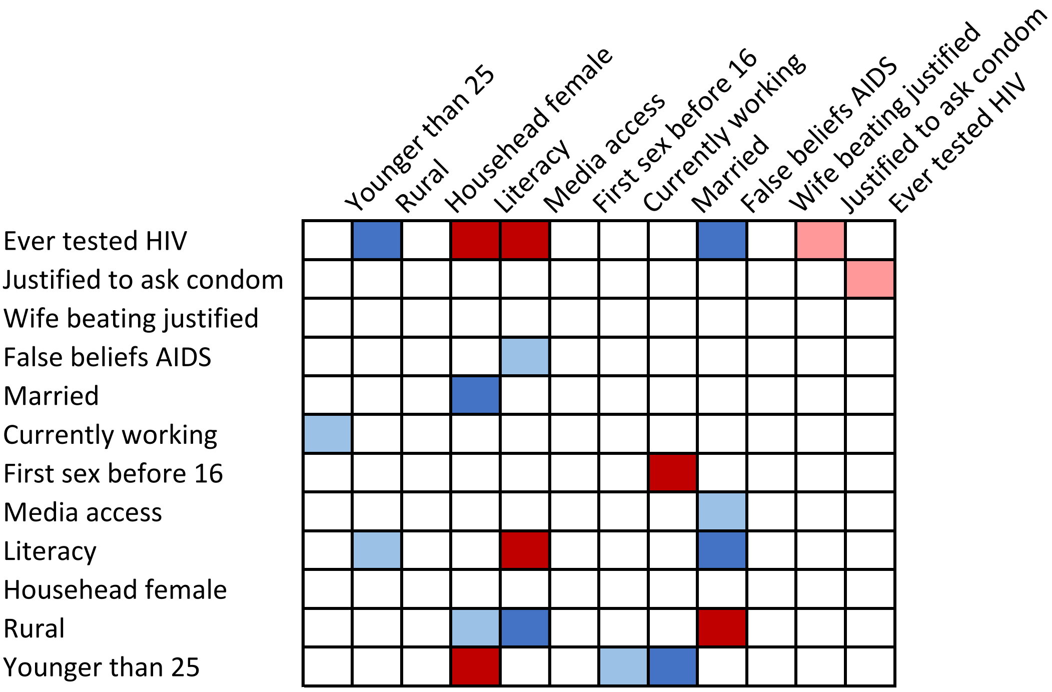


**Table S36. Mali, male**

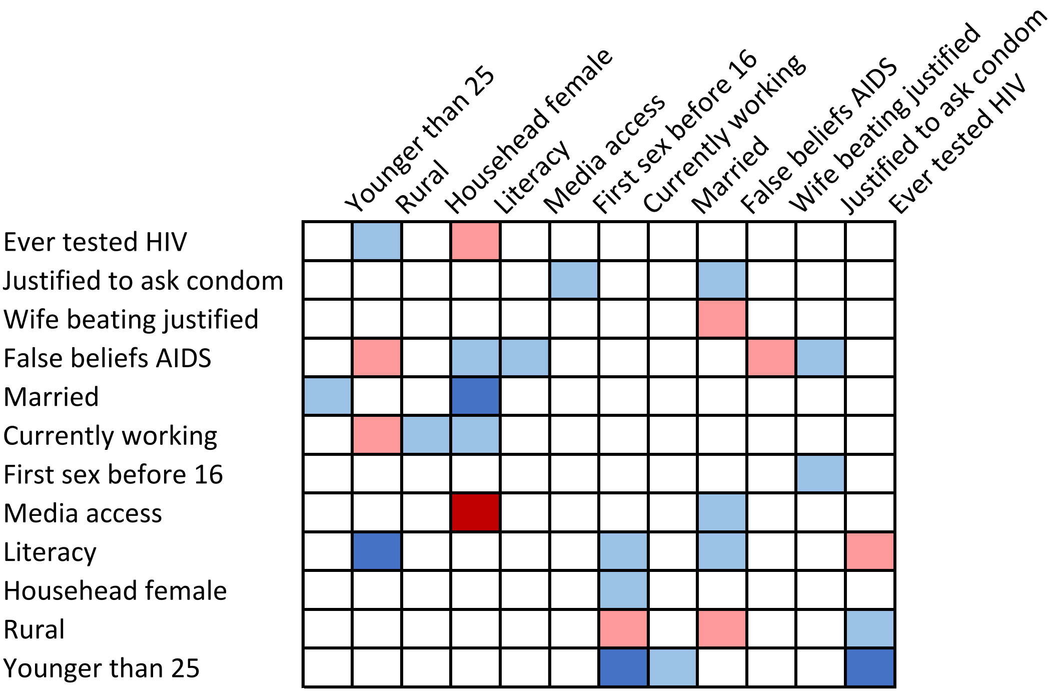

**Table S37. Mozambique, female**

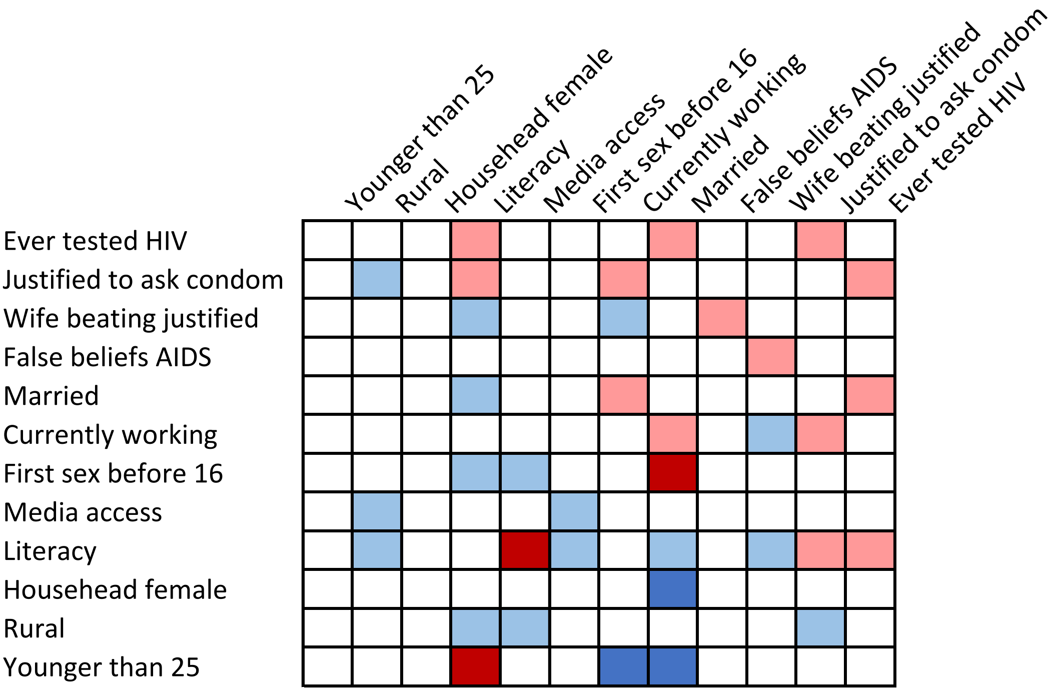


**Table S38. Mozambique, male**

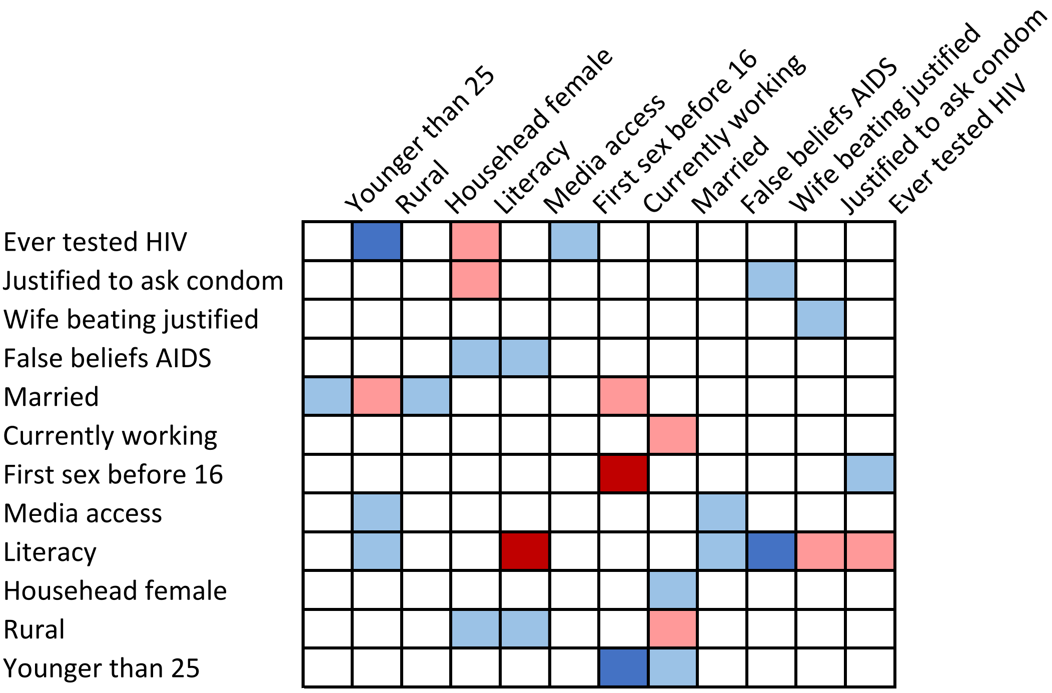

**Table S39. Namibia, female**

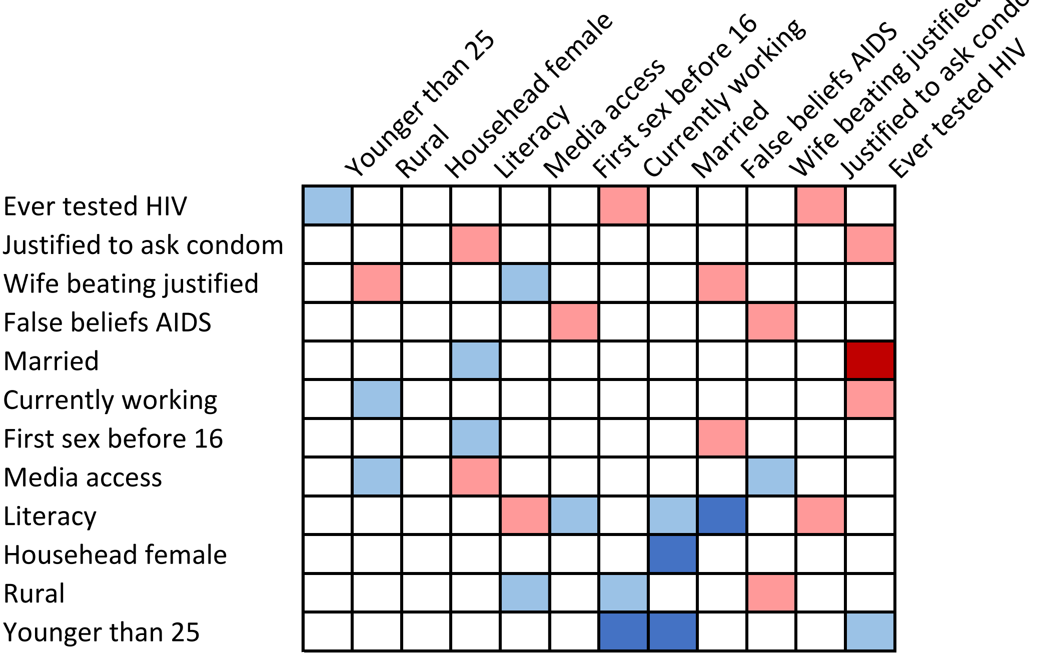


**Table S40. Namibia, male**

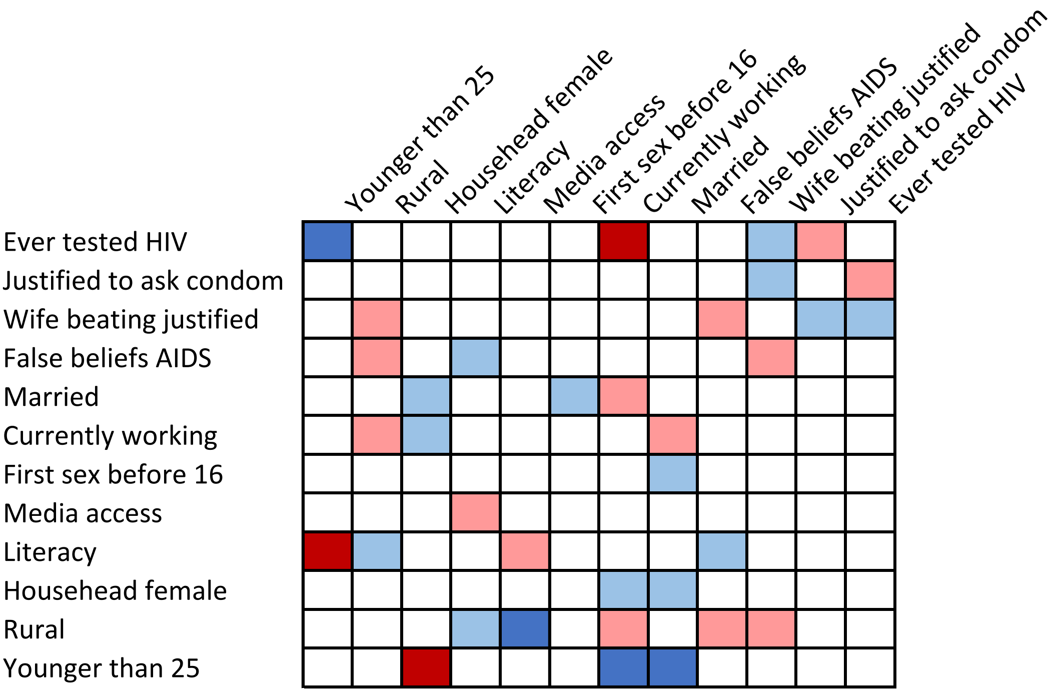

**Table S41. Niger, female**

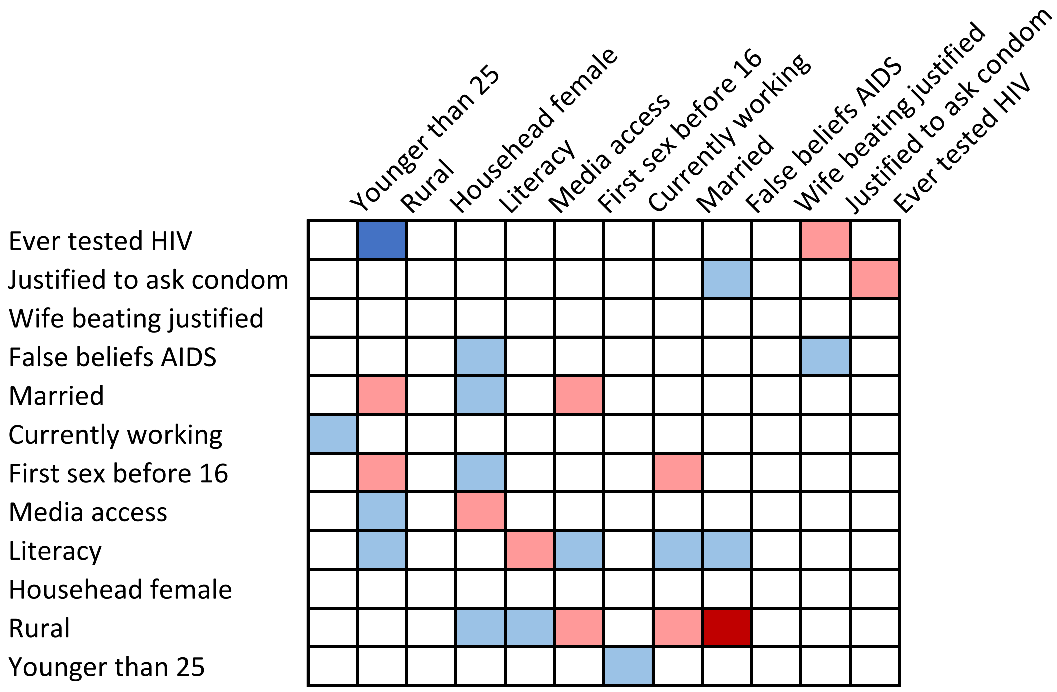


**Table S42. Niger, male**

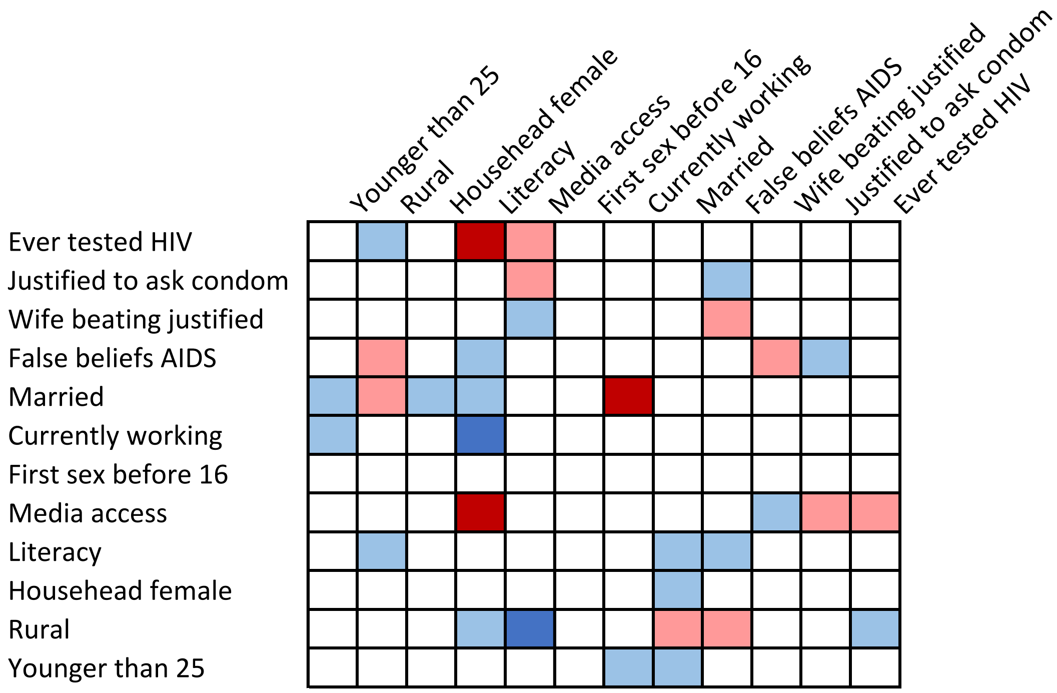

**Table S43. Nigeria, female**

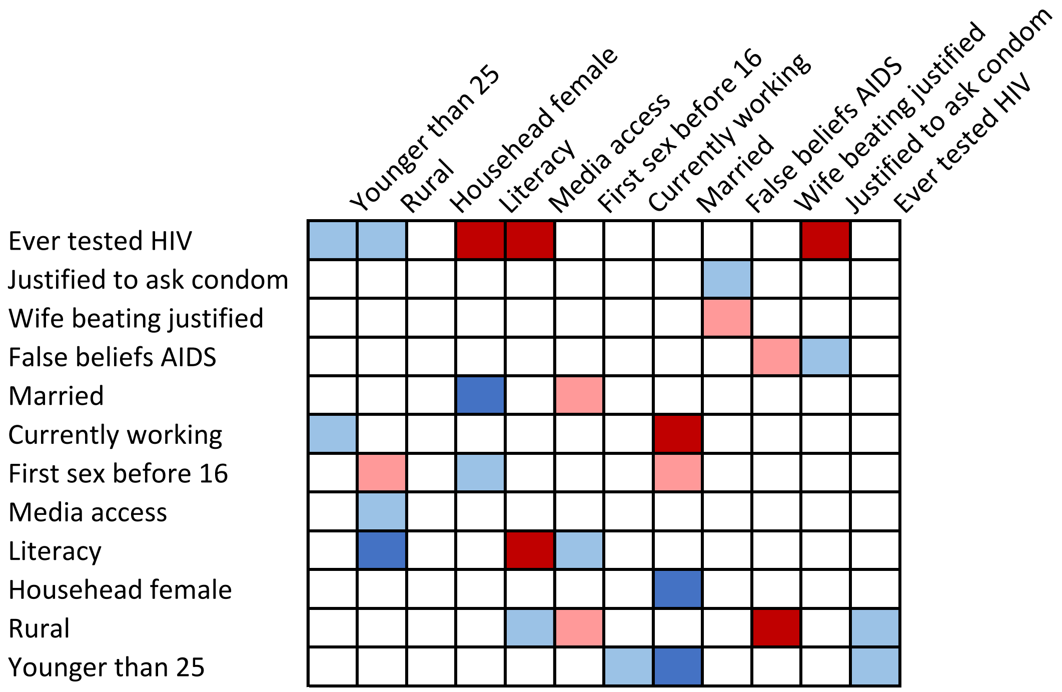


**Table S44. Nigeria, male**

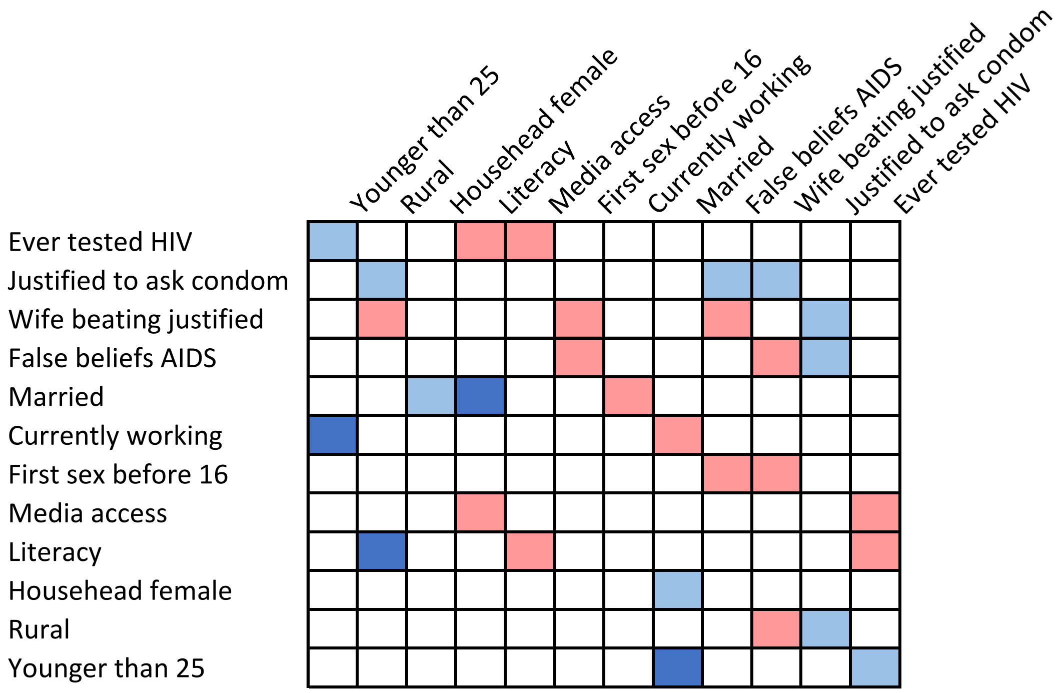

**Table S45. Rwanda, female**

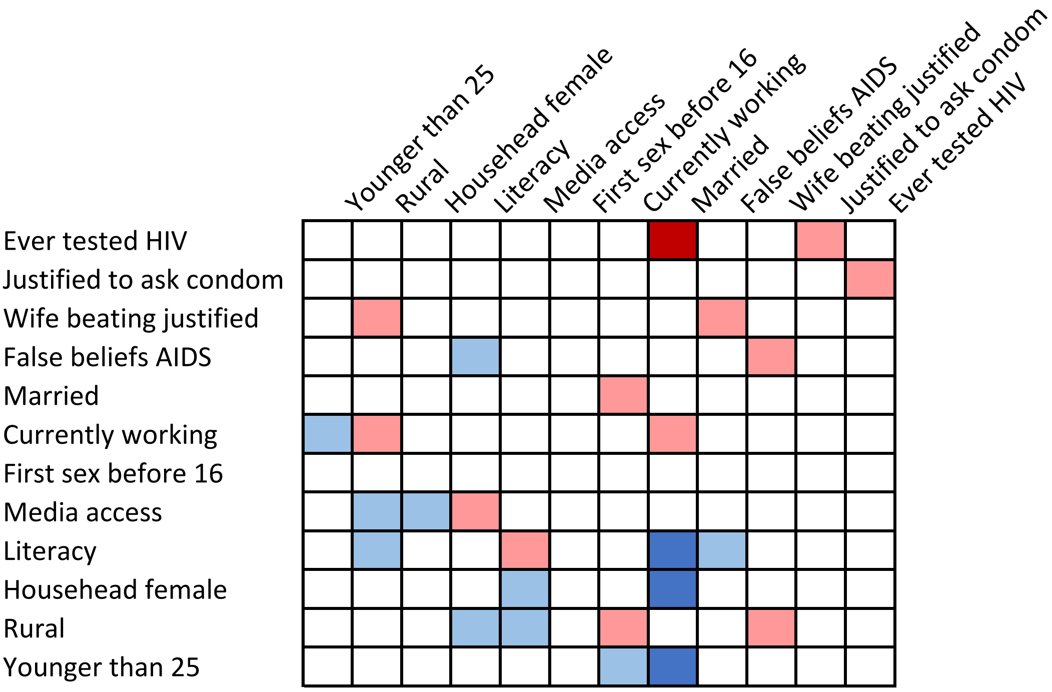


**Table S46. Rwanda, male**

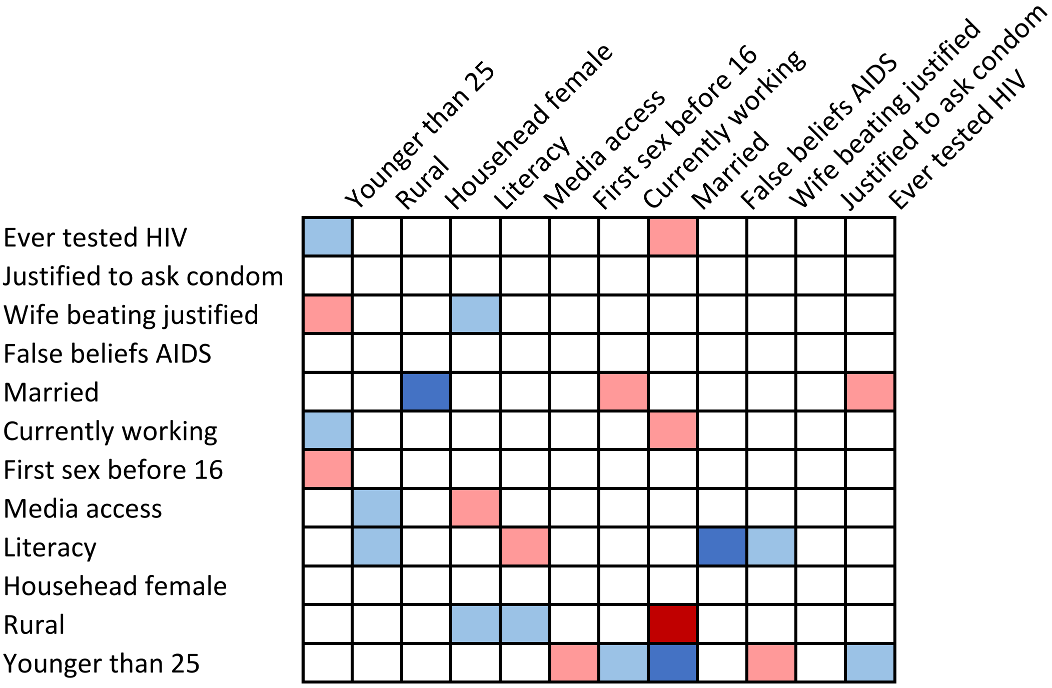

**Table S47. Senegal, female**

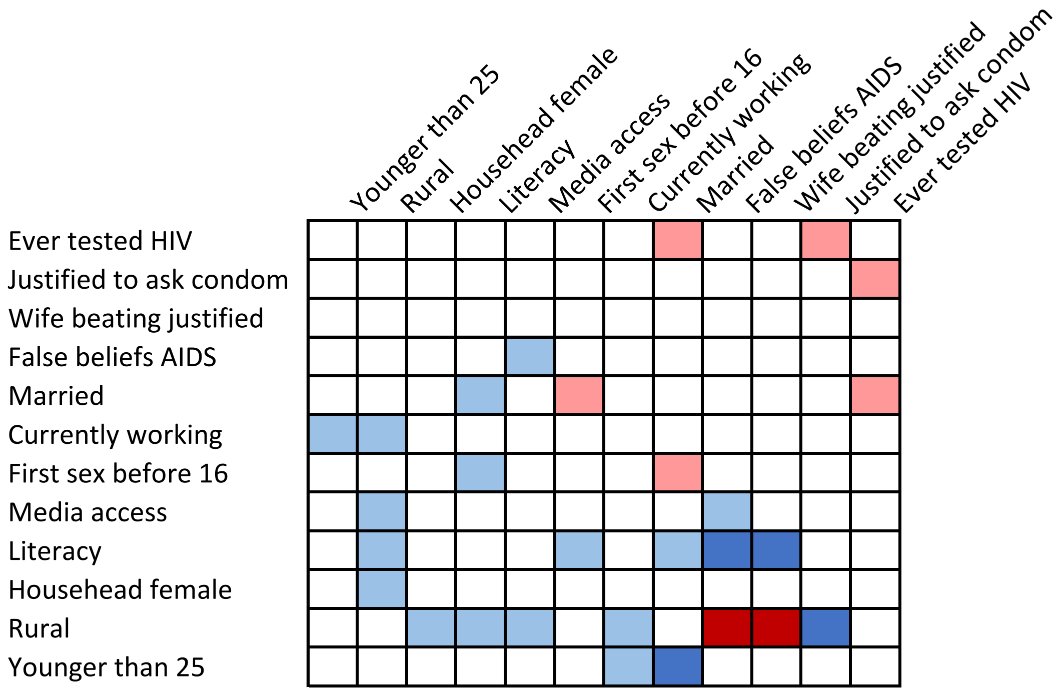


**Table S48. Senegal, male**

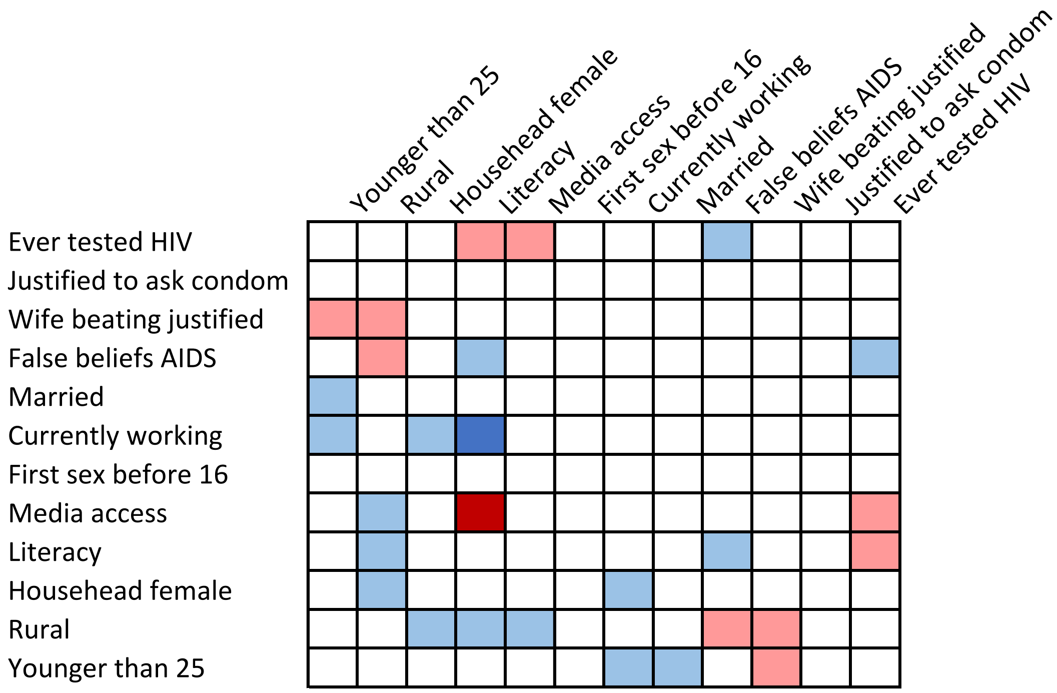

**Table S49. Sierra Leone, female**

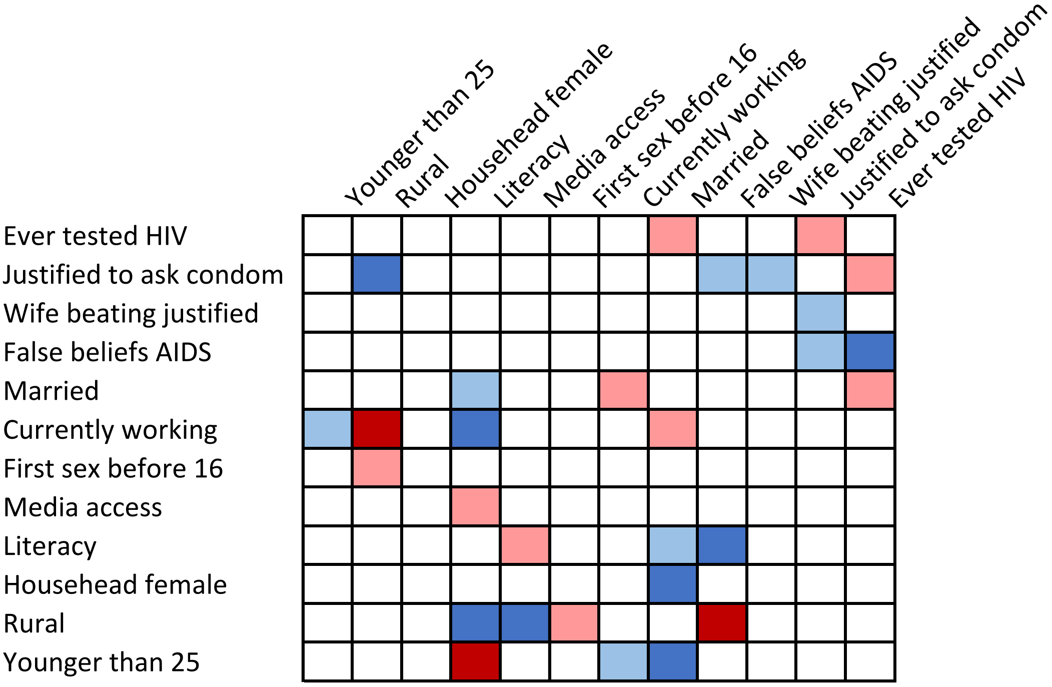


**Table S50. Sierra Leone, male**

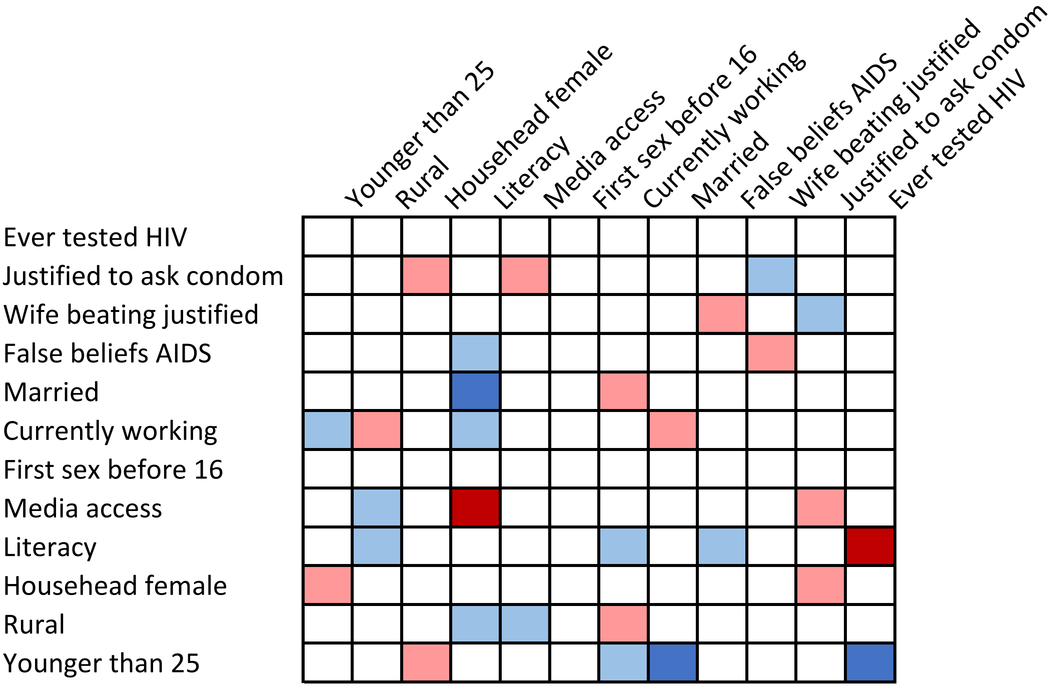

**Table S51. Togo, female**

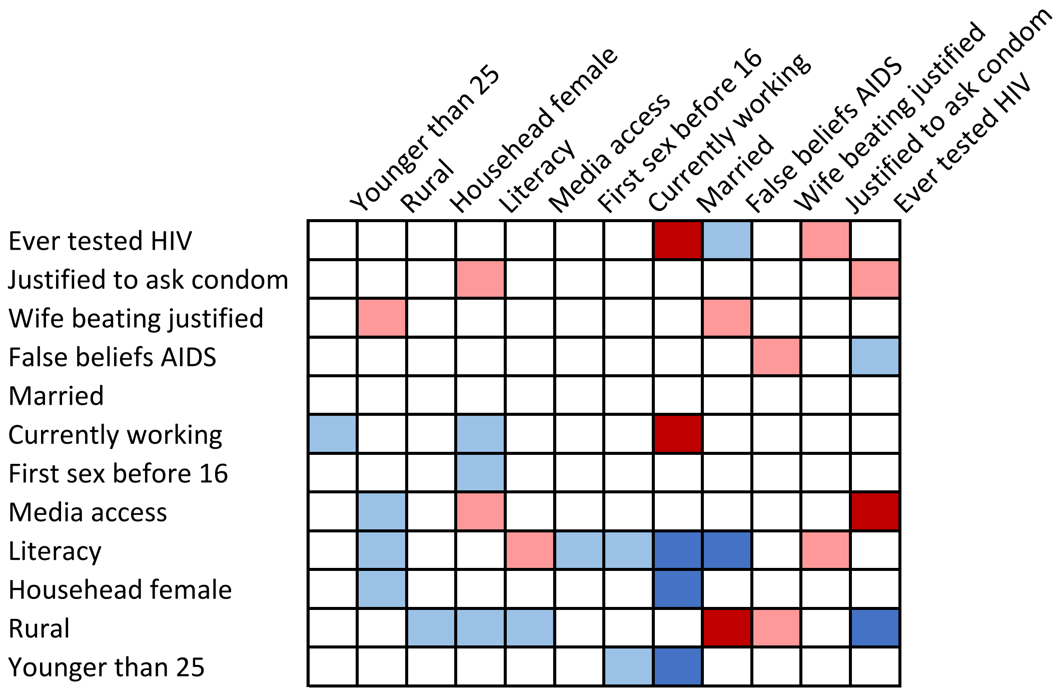


**Table S52. Togo, male**

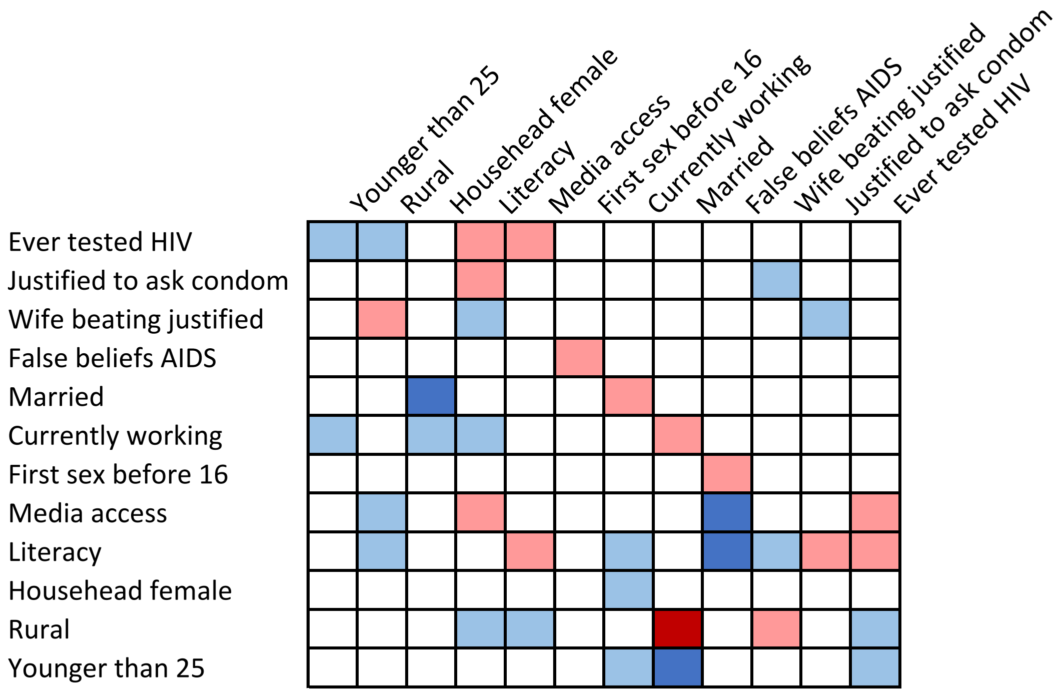

**Table S53. Uganda, female**

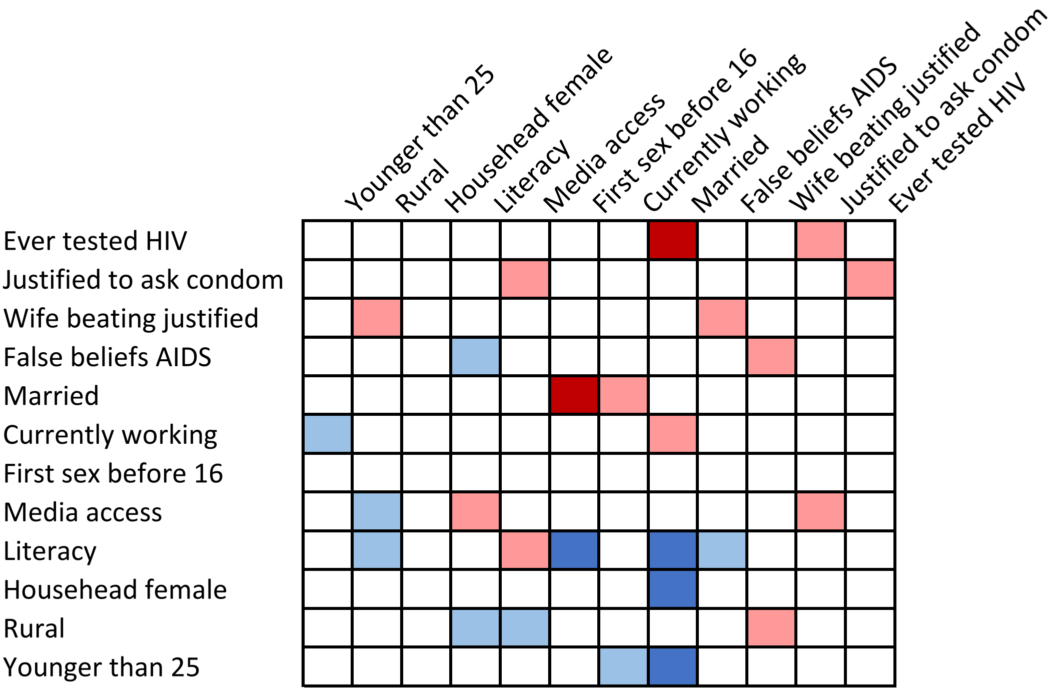


**Table S54. Uganda, male**

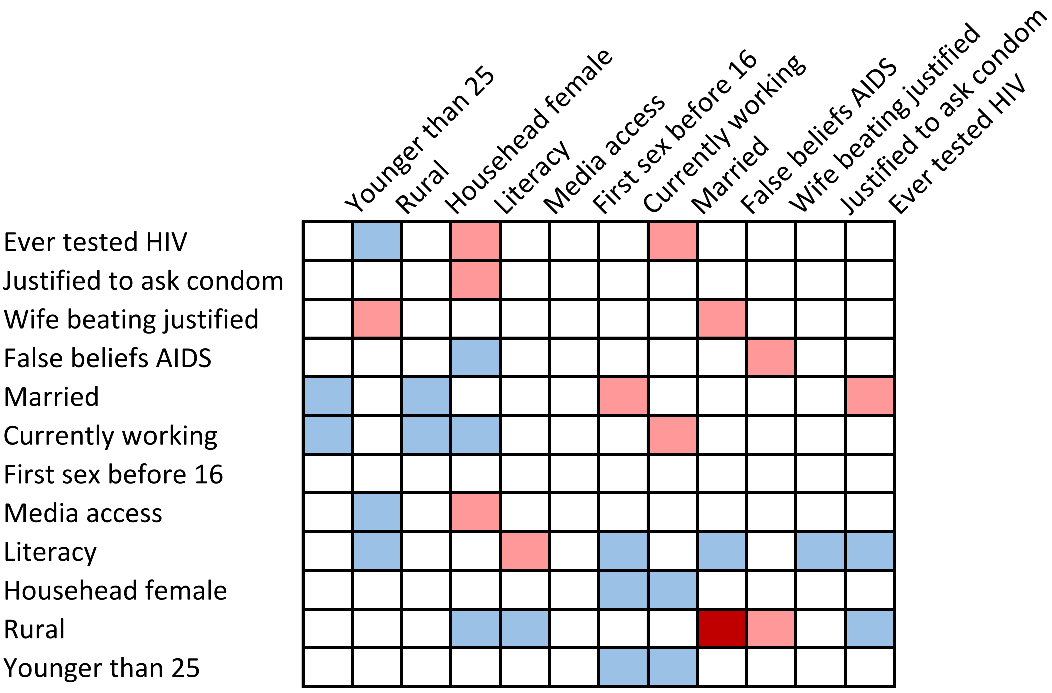

**Table S55. Zambia, female**

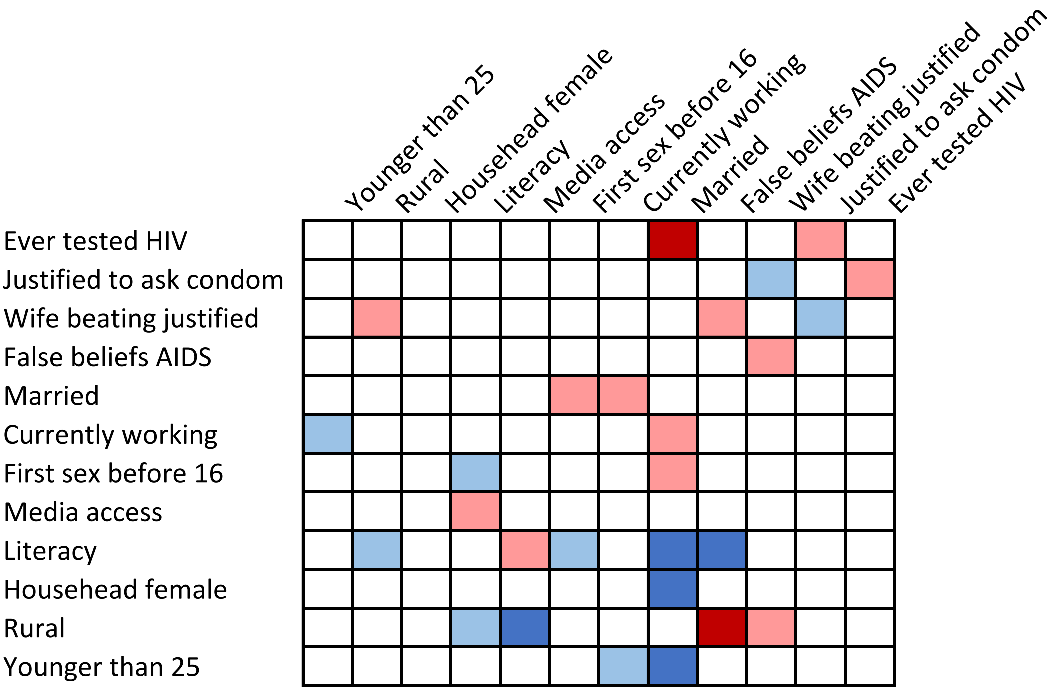


**Table S56. Zambia, male**

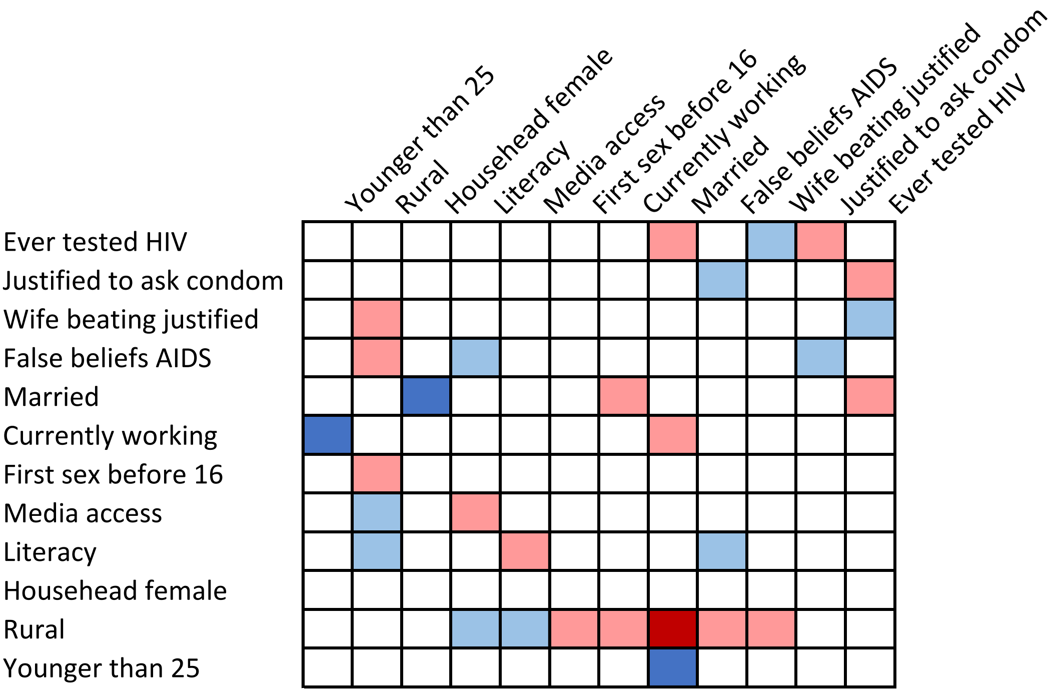

**Table S57. Zimbabwe, female**

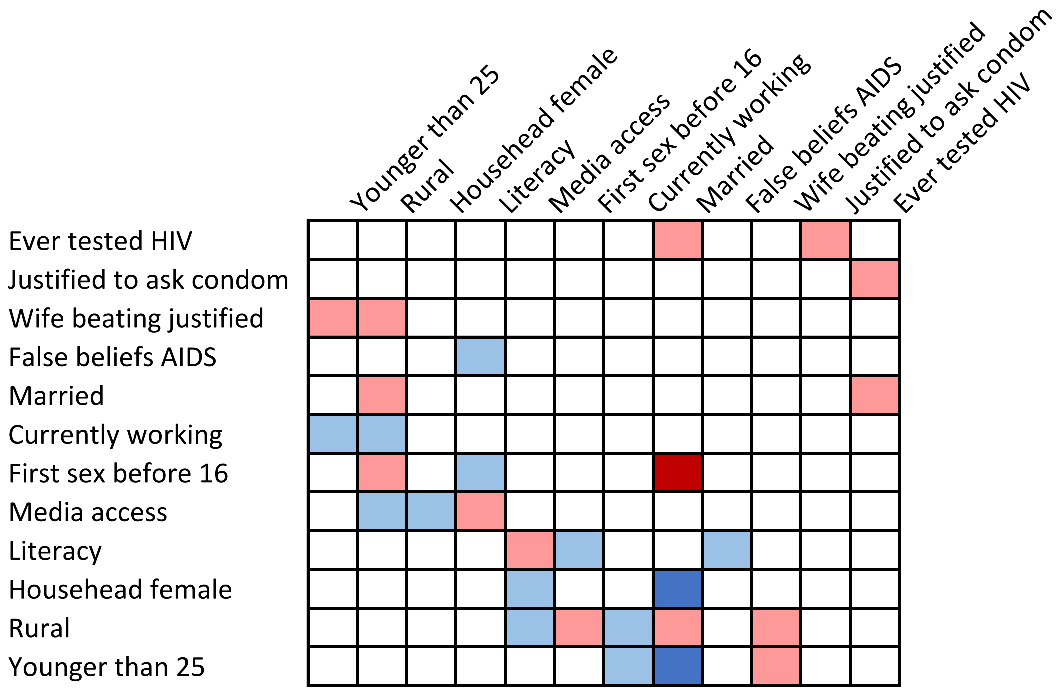


**Table S58. Zimbabwe, male**

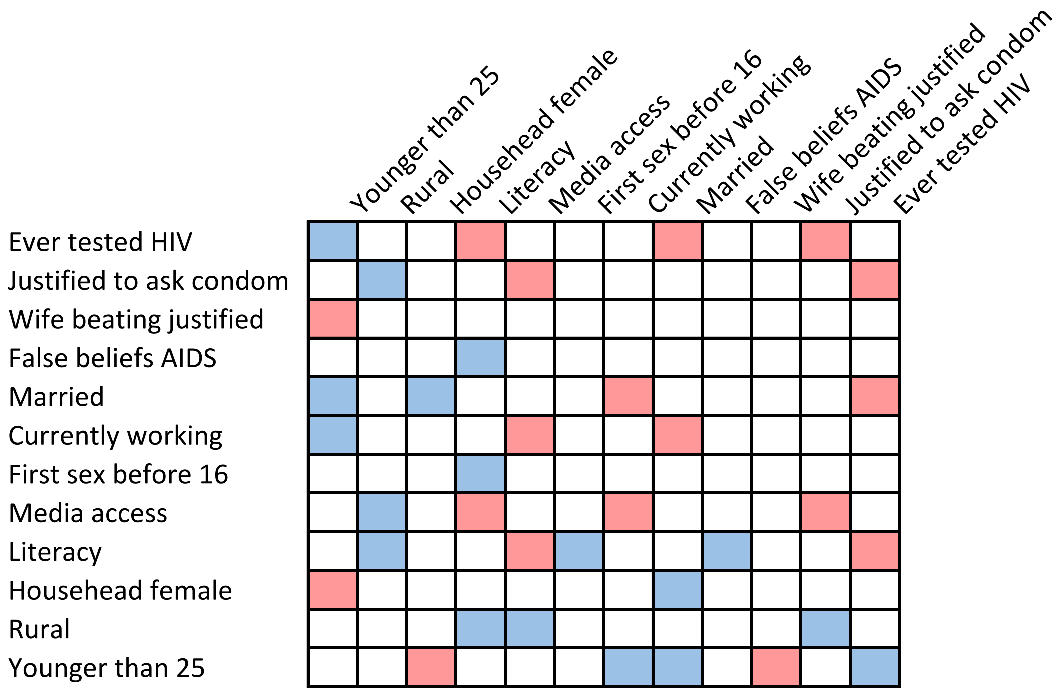
